# Evidence for both phylogenetic conservatism and lability in the evolution of secondary chemistry in a tropical angiosperm radiation

**DOI:** 10.1101/2020.11.30.404855

**Authors:** Kathryn A. Uckele, Joshua P. Jahner, Eric J. Tepe, Lora A. Richards, Lee A. Dyer, Kaitlin M. Ochsenrider, Casey S. Philbin, Massuo J. Kato, Lydia F. Yamaguchi, Matthew L. Forister, Angela M. Smilanich, Craig D. Dodson, Christopher S. Jeffrey, Thomas L. Parchman

## Abstract

Over evolutionary timescales, shifts in plant secondary chemistry may be associated with patterns of diversification in associated arthropods. Although foundational hypotheses of plant-insect codiversification and plant defense theory posit closely related plants should have similar chemical profiles, numerous studies have documented variation in the degree of phylogenetic signal, suggesting phytochemical evolution is more nuanced than initially assumed. We utilize proton nuclear magnetic resonance (^1^H NMR) data, chemical classification, and genotyping-by-sequencing to resolve evolutionary relationships and characterize the evolution of secondary chemistry in the Neotropical plant clade Radula (*Piper*; Piperaceae). Sequencing data substantially improved phylogenetic resolution relative to past studies, and spectroscopic characterization revealed the presence of 35 metabolite classes. Broad metabolite classes displayed strong phylogenetic signal, whereas the crude ^1^H NMR spectra featured evolutionary lability in chemical resonances. Evolutionary correlations were detected in two pairs of compound classes (flavonoids with chalcones; *p*-alkenyl phenols with kavalactones), where the gain or loss of a class was dependent on the other’s state. Overall, the evolution of secondary chemistry in Radula is characterized by strong phylogenetic signal of broad compound classes and concomitant evolutionary lability of specialized chemical motifs, consistent with both classic evolutionary hypotheses and recent examinations of phytochemical evolution in young lineages.

## Introduction

Plant secondary chemistry affects plant-herbivore interactions at various stages throughout an insect’s lifespan: mixtures of compounds can shape adult oviposition preferences (Thompson & Pellmyr, 1991), specific chemical compounds can stimulate larval feeding (Bowers, 1983, 1984), plant chemistry can deter insect herbivores via toxicity or physiological disruptions (Malcolm, 1994; Zagrobelny *et al*., 2004), and sequestered metabolites can alter immune function against natural enemies (Smilanich *et al*., 2009; Richards *et al*., 2012). Plants capable of developing novel chemical defenses are hypothesized to accrue higher fitness in response to enemy release (e.g., Berenbaum, 1978), potentially resulting in the diversification of plant lineages with conserved chemical phenotypes (the escape and radiate hypothesis; Ehrlich & Raven, 1964; Thompson, 1989; reviewed by Janz, 2011). Coevolutionary hypotheses and plant defense theory (reviewed by Mithöfer & Boland, 2012) have yielded clear predictions that herbivory, additional trophic interactions, and resource availability shape the evolution of plant defenses, including secondary metabolites (Agrawal *et al*., 2009; Maron *et al*., 2019). However, an evolutionary response to these biotic and abiotic pressures could be complex and highly context-dependent.

Due in part to the enzymatic complexity of metabolic biosynthesis, phylogenetic conservatism is the null hypothesis for the evolution of plant secondary chemistry (Agrawal & Fishbein, 2006; Salazar *et al*., 2018). Indeed, expectations of phylogenetic conservatism appear to hold at deep evolutionary scales; for example, the family Solanaceae is characterized by the presence of tropane alkaloids (Griffin & Lin, 2000), though they are consistently present in only 3 of 19 tribes (Datureae, Hyoscyameae, Mandragoreae) and sporadically found elsewhere (Wink, 2003). Further, recent work suggests more classes of secondary metabolites are phylogenetically conserved in large seed plant clades (e.g., eudicots and superasterids) than at lower taxonomic scales (e.g., orders and families) (Zhang *et al*., 2020). However, at shallower scales, numerous studies provide evidence for evolutionary lability in chemical traits within genera (e.g., Becerra, 1997; Kursar *et al*., 2009; Agrawal *et al*., 2009; Rasmann & Agrawal, 2011; Salazar *et al*., 2016; Moreira *et al*., 2018; Allevato *et al*., 2019), suggesting that surveys of phytochemical variation within young plant lineages might yield variable perspectives on the evolution of secondary chemistry. Adding further complexity, many studies have found evidence for strong evolutionary associations among chemical classes (Kariñho-Betancourt *et al*., 2015; Boachon *et al*., 2018; Allevato *et al*., 2019). For example, Johnson et al. (2014) found a strong positive correlation between flavonoids and phenolic diversity and a strong negative correlation between ellagitannins and flavonoids across a phylogeny of 26 evening primroses *(Oenethera:* Onagraceae). Such associations are relevant because they may reflect evolutionary constraints, and their causes may be varied. For example, positive associations may be associated with chemical defense syndromes (Agrawal & Fishbein, 2006; Agrawal, 2007) or synergistic effects of multiple classes on herbivore deterrence (Dyer *et al*., 2003; Richards *et al*., 2016). Alternatively, negative associations might be consistent with evolutionary tradeoffs or at least different optima in defense space (Agrawal, 2007; Johnson *et al*., 2014). By leveraging advances in organic chemistry and genomics, we stand to increase phylogenetic and metabolomic resolution to provide novel insight into the evolution of phytochemistry.

Recent advances in chemical ecology have improved perspectives on phytochemical diversity across a broad range of taxonomic groups and metabolite classes (Sedio, 2017; Dyer *et al*., 2018). High throughput processing of plant tissue, rapid advances in spectroscopy, and improved ordination and network analyses have enabled characterization of metabolomic variation across plant communities (Richards *et al*., 2016; Salazar *et al*., 2016, 2018; Dyer *et al*., 2018; Sedio *et al*., 2018; Ernst *et al*., 2019; Kang *et al*., 2019) and stand to enhance our understanding of phytochemical evolution across taxonomic scales (Sedio, 2017). Additionally, structural metabolomic approaches like ^1^H NMR can provide improved resolution of structural variation across a wide range of metabolite classes. Selection on the plant metabolome is inherently multivariate, arising from diverse herbivore communities and environmental conditions (Fine *et al*., 2006; Salazar *et al*., 2018), and even relatively small structural changes can impart disproportionate shifts in bioactivity. Thus, approaches that capture a larger proportion of the structural variation underlying phytochemical phenotypes could be well suited to addressing hypotheses concerning evolutionary patterns.

Next-generation sequencing data has reinvigorated phylogenetic analyses of traditionally challenging groups characterized by recent or rapid diversification (Wagner *et al*., 2013; Bagley *et al*., 2020; Léveillé-Bourret *et al*., 2020) as well as hybridization (Eaton & Ree, 2013; Carter *et al*., 2019; Hipp *et al*., 2020). Reduced representation DNA sequencing approaches [e.g., RADseq; genotyping-by-sequencing (GBS)] have been increasingly utilized in phylogenetic studies due to their ability to effectively sample large numbers of orthologous loci throughout the genomes of non-model organisms without the need for prior genomic resources (Leaché & Oaks, 2017; Parchman *et al*., 2018). Nearly all such studies have reported increased topological accuracy and support compared with past phylogenetic inference based on smaller numbers of Sanger-sequenced loci (Herrera & Shank, 2016; Massatti *et al*., 2016; Du *et al*., 2020), especially when applied to diverse radiations (Wagner *et al*., 2013; Fernández-Mazuecos *et al*., 2017; Hamon *et al*., 2017; Paetzold *et al*., 2019). While reduced representation approaches have clear phylogenetic utility at relatively shallow time scales, they have also performed well for moderately deep divergence (Eaton *et al*., 2017; Du *et al*., 2020).

*Piper* (Piperaceae) is a highly diverse, pantropical genus of nearly 2,600 accepted species (Callejas-Posada, 2020), with the highest diversity occurring in the Neotropics (Gentry, 1993; Martínez *et al*., 2015). Chemically, *Piper* is impressively diverse (Parmar *et al*., 1997; Dyer & Palmer, 2004; Richards *et al*., 2015): chemical profiling in a modest number of taxa has yielded 667 different compounds from 11 distinct structural classes thus far (Parmar *et al*., 1997; Dyer *et al*., 2004; Kato & Furlan, 2007; Richards *et al*., 2018). This phytochemical diversity has likely contributed to the diversification of several herbivorous insect lineages that specialize on *Piper*, including most notably the geometrid moth genus *Eois* (Strutzenberger *et al*., 2012; Wilson *et al*., 2012; Jahner *et al*., 2017). Furthermore, phytochemical variation in *Piper* communities has been shown to shape tri-trophic interactions and the structure of tropical communities (Dyer *et al*., 2004; Glassmire *et al*., 2016; Richards *et al*., 2018). As a species-rich genus with abundant and ecologically consequential phytochemical variation, *Piper* represents a valuable system for understanding how the history of diversification underlies the evolution of phytochemical variation.

*Piper* is an old lineage (~72 Ma), yet most of its diversification occurred in the Neotropics during the last 30-40 My following Andean uplift and the emergence of Central America (Smith *et al*., 2008; Martínez *et al*., 2015). The largest clade of *Piper*, Radula, exemplifies this pattern, as much of its extant diversity (~450 species) arose relatively recently during the Miocene (Martínez *et al*., 2015). Such bouts of rapid and recent diversification have limited the efficacy of traditional Sanger sequencing methods to resolve the timing and tempo of diversification in *Piper* (Jaramillo *et al*., 2008; Smith *et al*., 2008). Past phylogenetic analyses utilizing Sanger-sequenced nuclear and chloroplast regions have consistently inferred eleven major clades within *Piper*; however, phylogenetic resolution within these clades has been elusive (Jaramillo *et al*., 2008; Smith *et al*., 2008; Molina-Henao *et al*., 2016; Asmarayani, 2018). Phylogenetic inference based on genome-wide data spanning a range of genealogical histories has recently improved phylogenetic resolution for diverse radiations (e.g., Wagner *et al*., 2013; Paetzold *et al*., 2019), and should facilitate an understanding of evolutionary patterns of phytochemical variation in *Piper* and their consequences for plant-insect codiversification.

We leveraged complementary phylogenomic, metabolite classification, and ^1^H NMR data sets to generate a *Piper* phylogeny and explore the evolution of secondary chemistry within the largest *Piper* clade (Radula). Specifically, our goals were to: 1) resolve the evolutionary relationships within the Radula clade of *Piper* included in this study; 2) characterize metabolomic variation across the genus and within Radula in particular; and 3) quantify the strength of phylogenetic signal and detect evolutionary associations in Radula secondary chemistry. Because secondary chemistry is an emergent composite phenotype of many traits that can evolve semi-independently, we expected to detect mixed strengths of phylogenetic signal and strong associations among a subset of traits over evolutionary time.

## Materials and Methods

### Study system and sample collection

For phylogenetic and chemical analyses, we collected leaf material from 71 individuals representing 65 Neotropical *Piper* species from the following clades: Churumayu (*N* = 3), Hemipodium (*N* = 1), Isophyllon (*N* = 5), Macrostachys (*N* = 4), Peltobryon (*N* = 2), Pothomorphe (*N* = 1), Radula (*N* = 44), and Schilleria (*N* = 5). For chemical profiling and DNA sequencing, we collected the youngest, fully expanded leaves and dried them immediately with silica gel. Vouchers were pressed, dried, and deposited in one or more herbaria for future reference and species verification (Table S1). To investigate the evolution of phytochemistry at a relatively shallow evolutionary scale, we conducted the majority of our sampling within Radula (Martínez *et al*., 2015).

### Phylogenetic analyses

Genome-wide polymorphism data was generated for 71 individuals for phylogenetic analyses. Either the same accession sampled for chemical analysis, or an individual from the same population as the one sampled, were sequenced with a genotyping-by-sequencing approach (Parchman *et al*., 2012) that is analogous to ddRADseq (Peterson *et al*., 2012). Briefly, genomic DNA was digested with two restriction enzymes, *Eco*RI and *Mse*I. Sample-specific barcoded oligos containing Illumina adaptors were annealed to the *Eco*RI cut sites, and oligos containing the alternative Illumina adaptor were annealed to the *Mse*I cut sites. Fragments were PCR amplified and pooled for sequencing. The library was size-selected for fragments between 350 - 450 base pairs (bp) with the Pippin Prep System (Sage Sciences, Beverly, MA), and sequenced on two lanes of an Illumina HiSeq 2500 at the University of Texas Genome Sequencing and Analysis Facility (Austin, TX). Single-end, 100 bp, raw sequence data were filtered for contaminants (*E. coli*, *Phi*X, Illumina adaptors or primers) and low quality reads using `bowtie2_db` (Langmead & Salzberg, 2012) and a pipeline of bash and perl scripts (https://github.com/ncgr/tapioca). We used custom perl scripts to demultiplex our reads by individual and trim barcodes and restriction site-associated bases.

Assembly and initial filtering was conducted with `ipyRAD` v.0.7.30 (Eaton, 2014). `ipyRAD` was specifically designed to assemble RADseq data for phylogenetic applications, permits customization of clustering and filtering, and allows for indel variation among samples (Eaton, 2014). Because a suitable *Piper* genome was not available at the time of analysis, we generated a *de novo* consensus reference of sampled genomic regions with `ipyRAD`. Briefly, nucleotide sites with phred quality scores lower than 33 were treated as missing data. Sequences were clustered within individuals according to an 85% similarity threshold with `vsearch` (Rognes *et al*., 2016) and aligned with `muscle` (Edgar, 2004) to produce stacks of highly similar RADseq reads (hereafter, RADseq loci). The sequencing error rate and heterozygosity were jointly estimated for all RADseq loci with a depth >6, and these parameters informed statistical base calls according to a binomial model. Consensus sequences for each individual in the assembly were clustered once more, this time across individuals, and discarded if possessing >8 indels (max_Indels_locus), >50% heterozygous sites (max_shared_Hs_locus), or >20% variable sites (max_SNPs_locus). To reduce the amount of missing data in our alignment matrix, RADseq loci were retained if they were present in at least 50 of 71 samples. The nexus file of concatenated consensus sequences for each individual, including invariant sites, were used as input for the Bayesian phylogenetic methods described below. The nexus alignment as well as complete information on additional parameter settings for this analysis are archived at Dryad (https://doi.org/10.5061/dryad.j6q573nc7).

To resolve patterns of diversification and to provide a foundation for investigating variation in the rates of phytochemical evolution, we estimated a rooted, calibrated tree according to a relaxed clock model in `RevBayes` v.1.0.12 (Höhna *et al*., 2016), which provides the ability to specify custom phylogenetic models for improved flexibility compared with other Bayesian approaches. The prior distribution on node ages was defined by a birth-death process in which the hyper priors on speciation and extinction rates were exponentially distributed with *λ* = 10. We relaxed the assumption of a global molecular clock by allowing each branch-rate variable to be drawn from a lognormal distribution. After comparing the relative fits of JC, HKY, GTR, and GTR+Gamma nucleotide substitution models with Bayes factors, we modeled DNA sequence evolution according to the best-fit HKY model. Eight independent MCMC chains were run for 100,000 generations with a burn-in of 1,000 generations and sampled every 10 generations. Chains were visually assessed for convergence with `Tracer` v.1.7.1 (Rambaut *et al*., 2018) and numerically assessed with effective sample sizes (ESS), the Gelman-Rubin convergence diagnostic (Gelman & Rubin, 1992), and by comparing the posterior probabilities of clades sampled between MCMC chains. The maximum clade credibility (MCC) tree provided the ultrametric fixed tree topology and relative node ages for phylogenetic comparative methods described below.

### Chemical profiling

Crude proton nuclear magnetic resonance (^1^H NMR) spectroscopy was chosen for chemotype mapping due to its ability to characterize subtle structural variation across a wide range of compound classes in a single, reproducible, non-destructive analysis (Richards *et al*., 2018). Briefly, after leaf samples were ground to fine powder, 2.00 g were transferred to a glass screw cap test tube with 10.0 ml of methanol, sonicated for 10 minutes, and filtered. This step was repeated and both filtrates were combined in a pre-weighed 20 ml scintillation vial. The solvent was removed *in vacuuo* and dissolved in 0.6 ml methanol-*d*_4_ for ^1^H NMR analysis. Extracts were analyzed on a Varian 400 MHz solution state NMR spectrometer with autosampler. Data were processed using MestReNova software (Mestrelab Research, Santiago de Compostela, Spain). Spectra from the crude extracts were aligned with the solvent peak (CD_3_, δ = 3.31 ppm), baseline corrected, phase corrected, and binned (0.04 ppm; 0.5 - 12 ppm). Solvent and water peaks were removed and the binned spectra were normalized to a total area of 100. This data set is referred to subsequently as “crude ^1^H NMR”.

In addition to crude ^1^H NMR spectral chemotyping, we further annotated and characterized samples based upon the presence or absence of compound classes and in some cases, specific compounds. To further gain structural resolution across the crude extracts that were sampled, aliquots of the ^1^H NMR extracts were diluted and subjected to GC-MS and LC-MS analysis. Crude extracts were classified using chemotaxonomic classifications outlined in Parmar’s comprehensive review of *Piper* phytochemistry (Parmar *et al*., 1997).

Presumptive compounds and compound classes were annotated based upon structural elucidation using ^1^H NMR, GC-MS fragmentation, and high-resolution LC-MS data. Comparison of the ^1^H NMR data to literature values of related compounds was used to increase confidence in these assignments. In some cases, crude 2D-NMR analysis was used to confirm structural classifications. Presence of a compound or compound class was determined based upon abundant and spectroscopically apparent evidence. This data set is referred to subsequently as “metabolite classes”.

### Phylogenetic signal and evolution of metabolite classes

To assess whether metabolite classes were phylogenetically conserved across Radula, we quantified phylogenetic signal in these binary traits using the D statistic (Fritz & Purvis, 2010). The D statistic calculates the sum of sister-clade differences, ∑d_obs_ (Felsenstein, 1985) for an observed tree and binary trait, and scales this value with the distributions of sums expected under two disparate evolutionary models, random and Brownian motion (∑d_r_ and ∑d_b_, respectively), using the following equation:

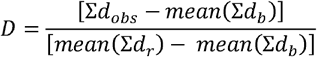

Thus, D is expected to equal 1 when the observed binary trait is distributed randomly, lacking phylogenetic signal, and is expected to equal 0 when it is exhibits phylogenetic signal as expected under Brownian motion. Tests of phylogenetic signal with the D statistic are most accurate when the ratio of presences and absences is closer to 1:1 (Fritz and Purvis, 2010). We used the *phylo.d* function in the `caper` package (Orme *et al*., 2018) in `R` v.4.0.0 (R Core Team, 2020) to calculate the observed D for a subset of binary traits that were sufficiently present across the phylogeny. This value was compared to a distribution of D values simulated under models of phylogenetic randomness (D = 1) and pure Brownian motion (D = 0) to determine whether the observed D differed from either zero or one.

To detect evolutionary associations among pairs of metabolite classes within Radula, we used Pagel’s (1994) method that models evolutionary changes in two binary traits, X and Y, as continuous-time Markov processes in which the probabilities of state transition at one trait may depend on the state at the other trait. Significant tests of correlated evolution were followed by tests of contingency, in which changes at X depend on the state of Y, or vice versa. Model fits, comparisons, and plots were performed with the *fitPagel* function in the `phytools` package (Revell, 2012) in `R`.

### Multivariate analyses of phylogenetic signal with crude ^1^H NMR spectra

While the analyses above based on broad classifications of structurally determined metabolites provide a coarse view of phytochemical evolution, these classifications are anchored to the foundations of plant secondary metabolite biosynthesis. Using ^1^H NMR spectra as a raw chemotype should allow a more detailed multivariate perspective on phytochemical variation. Studies on other plant taxa have typically detected some signal and evolutionary correlations for broad classes of compounds but not necessarily for specific compounds or biologically active moieties, both of which can be inferred from ^1^H NMR data. Multivariate approaches to phylogenetic comparative methods have provided insight into covarying suites of related traits, while simultaneously increasing the statistical power to detect phylogenetic signal (Zheng *et al*., 2009) and differences in trait means among taxa (Clavel *et al*., 2015). Indeed, these multivariate approaches might be particularly useful when exploring the evolution of complex phenotypes, like the plant metabolome, which exhibit trait covariances due to metabolomic or functional associations (Dyer *et al*., 2003; Richards *et al*., 2010; Fukushima *et al*., 2011). Here we utilize three multivariate methods to detect patterns of phylogenetic signal for 263 resonances found in the crude ^1^H NMR data: 1) principal components analyses (PCA); 2) multiple regression on distance matrices (MRM); and 3) multivariate estimation of phylogenetic signal.

To visualize patterns of chemotypic variation across all sampled species from all clades, we first analyzed the ^1^H NMR data with PCA using the *prcomp* function in `R`. If the major axes of metabolomic variation are phylogenetically conserved, the plotted species scores should be clustered by clade in a rotated principal component (PC) space. Alternatively, if metabolomic variation is randomly distributed across the phylogeny, there should be little to no clustering by clade (Klingenberg & Gidaszewski, 2010). The degree to which plant clade predicted chemical similarity was assessed using permutational multivariate analysis of variance (permanova; Anderson, 2001) in the `vegan` package (Oksanen *et al*., 2019) in R based on Euclidean distances of the first four PCs.

Mantel tests have been frequently used to assess the degree of phylogenetic signal in multivariate data (e.g., Cardini & Elton, 2008; Easson & Thacker, 2014; Salazar *et al*., 2018) by estimating the relationship between phylogenetic and phenotypic distances. Simulations under scenarios of measurement error have found instances where Mantel tests outperform traditional univariate methods in detecting phylogenetic signal, especially as the number of traits increases (Hardy & Pavoine, 2012). Because we were unable to account for measurement error in our study, we utilized MRM to examine the relationship between metabolomic and phylogenetic distance at two evolutionary scales (within Radula and across all clades). Euclidean distances were calculated with the crude ^1^H NMR spectra using the *dist* function in `R`, and two measures of phylogenetic distance were used as predictors: 1) Abouheifs proximity (Abouheif, 1999; Pavoine *et al*., 2008) was calculated using the *proxTips* function in the `adephylo` package (Jombart *et al*., 2010) in `R`; and 2) the square root of patristic distance was calculated using the *cophenetic.phylo* function in the `ape` package (Paradis *et al*., 2004) in `R`. MRM analyses were implemented using the *MRM* function with 1000 permutations in the `ecodist` package (Goslee & Urban, 2007) in `R`.

Since Blomberg et al.’s (2003) *K* statistic exhibits higher statistical power to detect phylogenetic signal relative to Mantel tests (Harmon & Glor, 2010), we quantified phylogenetic signal of the crude ^1^H NMR at both evolutionary scales using a multivariate generalization of the *K* statistic (*K*_mult_; Adams, 2014) with the *physignal* function in the `geomorph` package (Adams *et al*., 2013) in `R`. The *K* statistic provides a statistical estimate of phylogenetic signal relative to expectations under Brownian motion, where values of *K* greater than 1 indicate phylogenetic signal greater than expected under Brownian motion, whereas values between 0 and 1 indicate less signal than expected under Brownian motion. Significance for the generalized *K* statistic was assessed by permuting the ^1^H NMR peak data among the tips of the phylogeny for 999 iterations.

To determine whether the zero-inflated nature of the ^1^H NMR data influenced the detection of phylogenetic signal, we permuted our ^1^H NMR data set over 1000 iterations by randomly indexing our original ^1^H NMR data matrix. This permutation method preserves the original proportion of zeros in the matrix while obfuscating any observed phylogenetic signal. The generalized *K* statistic test was calculated for each permutation, and our observed generalized *K* statistic was compared to the null distribution of permuted values.

## Results

### Phylogenetic analyses

After contaminant filtering and demultiplexing, we retained ~313 million Illumina reads for phylogenetic analyses. Initial clustering, variant calling, and filtering clustered reads into 362,169 RADseq loci. There was a high proportion of missing data, presumably due to allelic dropout increasing with high levels of divergence among *Piper* clades. For Bayesian phylogenetic inference, we mitigated the influence of missing data by removing loci absent in >30% of samples. The final dataset for phylogenetic analyses consisted of 641 RADseq loci (~86 bp in length each) that housed 9,113 genetic variants (51% parsimony informative). Aligned loci were concatenated into a nexus alignment with missing data at 18.9% of sites.

Bayesian phylogenetic analysis of ddRADseq data resolved eight major Neotropical *Piper* clades with high posterior support (Fig. 1). While past phylogenetic studies supported the monophyly of seven of these eight clades (Macrostachys, Radula, Peltobryon, Pothomorphe, Hemipodion, Isophyllon, and Schilleria) (Jaramillo *et al*., 2008; Martínez *et al*., 2015), our analysis resolved an additional clade, Churumayu. Notably, Isophyllon and Churumayu were highly supported, monophyletic clades and not nested within Radula as was inferred in previous analyses (Jaramillo *et al*., 2008). Contrary to previous phylogenetic hypotheses of *Piper* Jaramillo *et al*., 2008; Martínez *et al*., 2015), our analyses might suggest Churumayu is the most basal clade, but we caution that this node had very low posterior support (51%). Intrageneric relationships below the clade level were highly resolved, with nearly all nodes exhibiting greater than 95% posterior support (Fig. 1), including within the diverse Radula clade (Fig. 1). Our phylogenetic hypothesis for Radula indicates three species (*P. hispidum, P. colonense, P. lucigaudens*) may be paraphyletic, reflecting past taxonomic uncertainty for these taxa.

**Figure 1.**
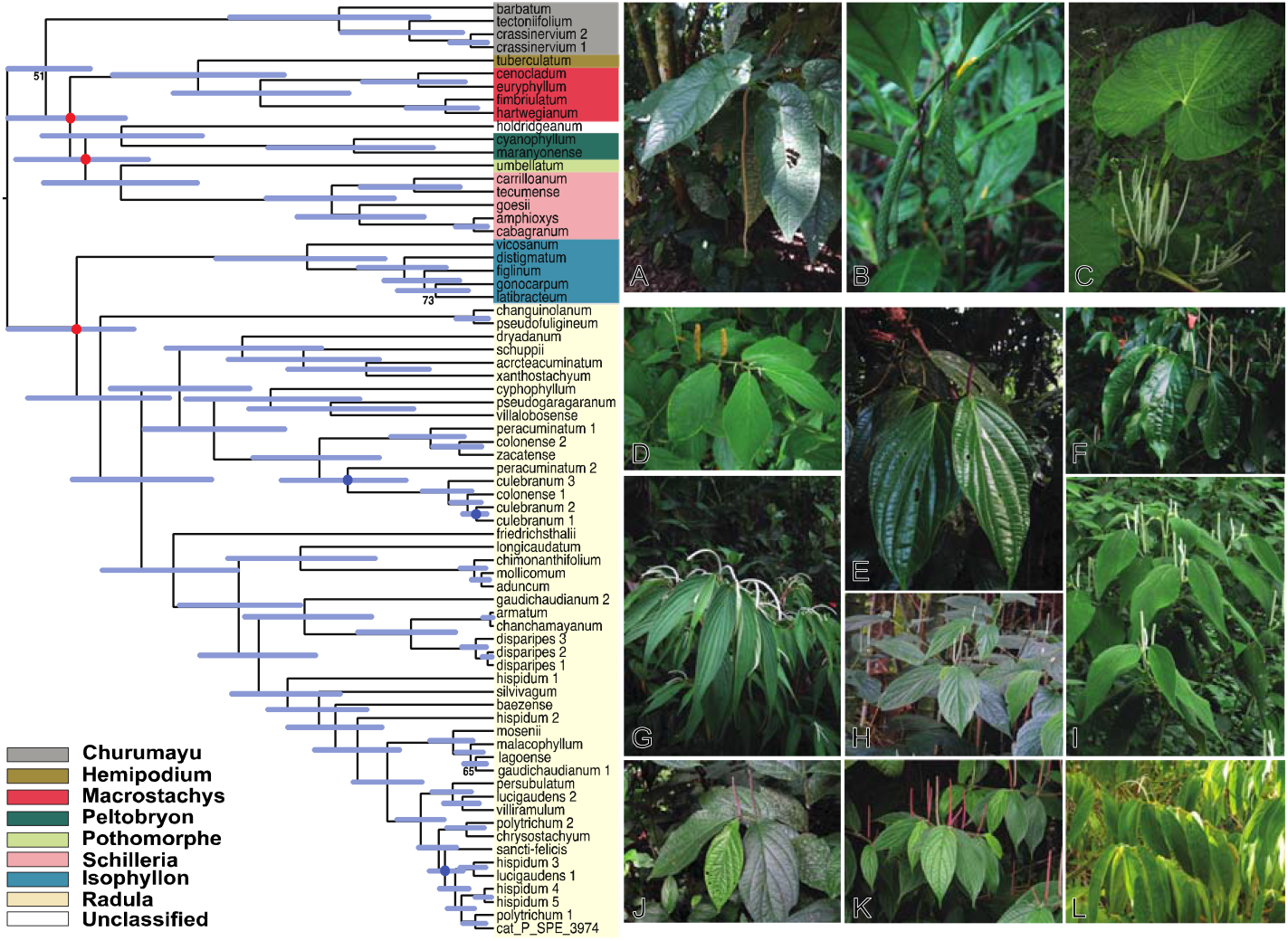
Maximum clade credibility tree of 48 species from the Radula clade of *Piper* and 23 outgroup species inferred with a Bayesian analysis of 641 concatenated RADseq loci (55,298 base pairs) comprising 9,113 genetic variants (of which 4,674 are parsimony informative). The outgroup taxa were sampled across multiple *Piper* clades: Isophyllon, Churumayu, Macrostachys, Hemipodium, Peltobryon, Pothomorphe, and Schilleria. All nodes are supported by at least 95% posterior support except where noted with circles or labels. Blue circles indicate support values between 85-95%. Red circles indicate support values between 75-85%. Three nodes with less than 75% posterior support were given numerical support values. Blue bars at each node denote the 95% highest posterior density interval on node ages. Diversity of *Piper* with the clade they belong to in parentheses. Images of outgroups include **A.** *Piper hillianum* (Macrostachys), **B.** *P. acutifolium* (Peltobryon), and **C.** *P. umbellatum* (Pothomorphe). Examples of the Radula clade of *Piper* include **D.** *P. pseudofuligineum*, **E.** *P. concepcionis*, **F.** *P. disparipes*, **G.** *P. friedrichsthalii*, **H.** *P. dilatatum*, **I.** *P. bredemeyeri*, **J.** *P. immutatum*, **K.** *P. erubescentispicum*, and **L.** the widespread and often weedy *P. aduncum*.

### Phytochemical variation in *Piper*

Nearly all common compound classes that have been previously reported in *Piper* were observed from our compound characterization analysis (Salehi *et al*., 2019). This analysis revealed the presence of metabolite classes that are ubiquitous across plant families (lignans, flavonoids/chalcones, etc.) as well as classes that are specifically common in *Piper* (amides) (Fig. 2). Specific compound characterization revealed genus specific compounds and compound classes (piplartine, cenocladamide, crassinervic acid, kava lactones), as well as metabolites that are more rarely reported in plants (putrescine diamides, nerolidyl catechol, alkenyl phenols, anuramide peptides) (Fig. 2).

**Figure 2.**
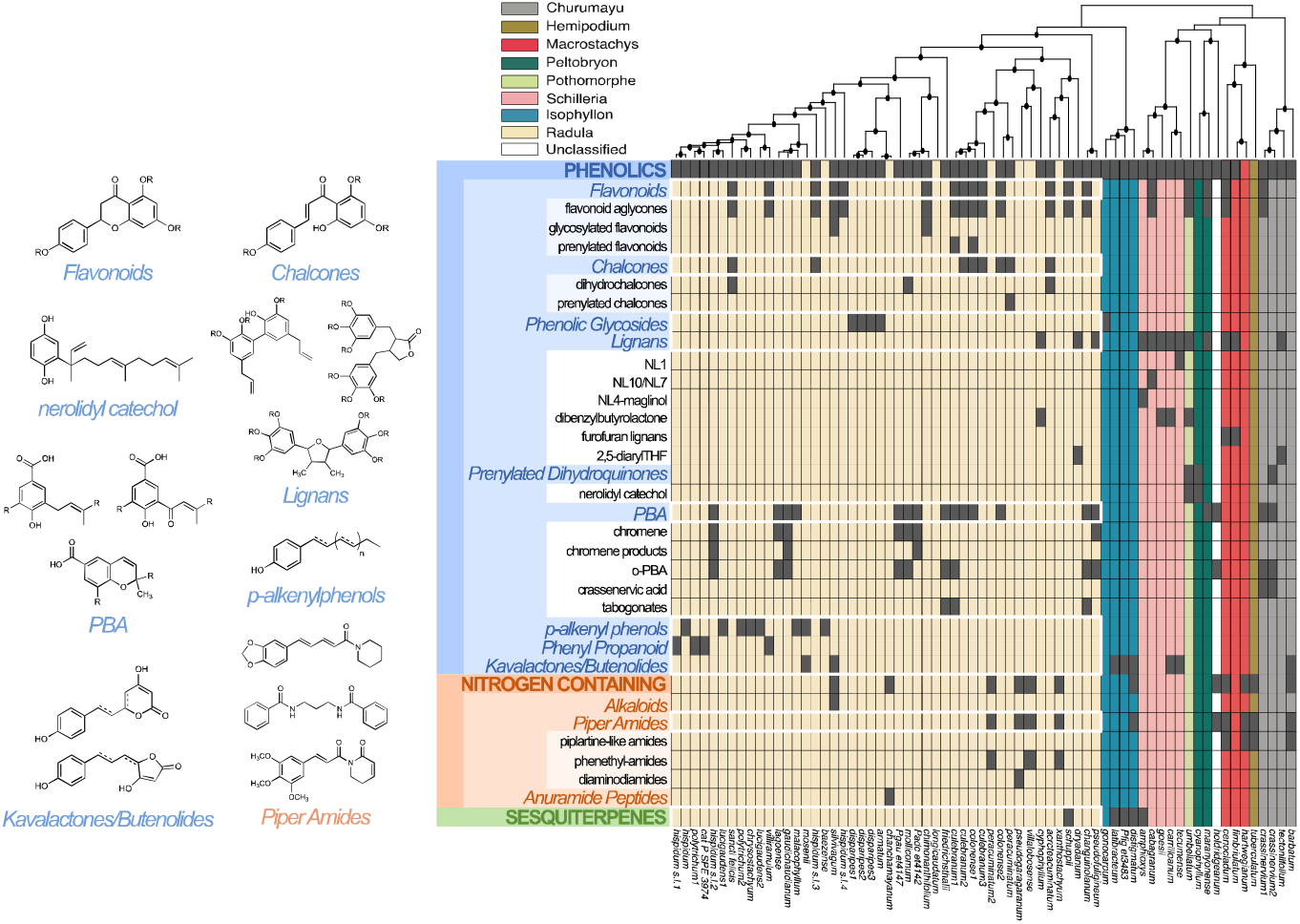
Taxa comprise the columns of the matrix and are ordered according to their inferred phylogenetic relationships. Groups of columns are colored according to their designated *Piper* clade. Black circles within the phylogenetic tree designate nodes with posterior support values greater than 85%. Each row of the matrix represents a metabolite class which was detected from ^1^H NMR, GC-MS, and LC-MS data, with dark grey cells indicating the presence of that class in that taxa. Rows outlined in white indicate traits which were analyzed for phylogenetic signal in Radula. To the left of the matrix are representative compounds for a subset of metabolite classes which were detected in our samples.

### Metabolite phylogenetic signal and evolutionary associations

For all eight metabolite classes that were examined, estimates of D (Fritz & Purvis, 2010) were low and did not deviate from a null distribution generated under a scenario of Brownian motion (Table 1), consistent with phylogenetic signal. Two of the eight traits, phenolic glycosides and lignans, exhibited strong phylogenetic signal (D < 0), while the remaining six traits exhibited weak phylogenetic signal (0 < D < 1). Further, all metabolite classes had observed values of D that differed from a null distribution generated under a phylogenetic randomness scenario (Table 1). The mean of the observed D estimates for the metabolite classes was 0.04, with the largest D statistic observed for the flavonoid class (*d_obs_* = 0.49) and the smallest observed for the phenolic glycosides (*d_obs_* = −1.18) (Table 1).

**Table 1.**
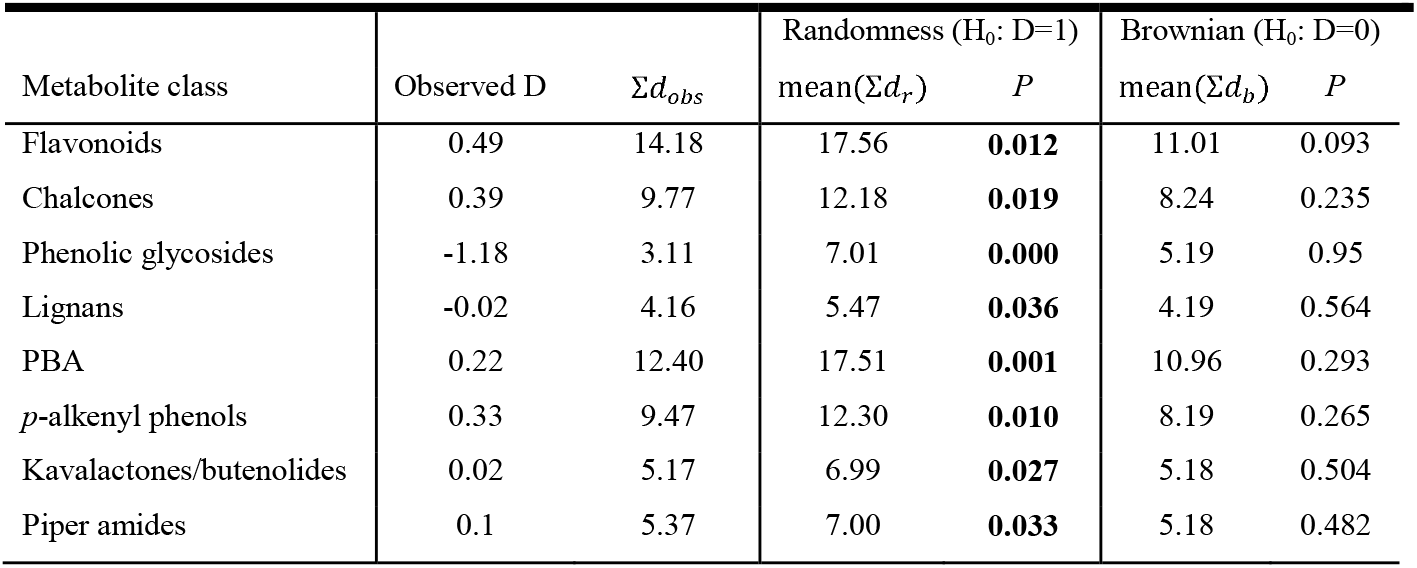
Estimates of phylogenetic signal (D) (Purvis and Fritz, 2010) for a subset of metabolite classes (see Methods for explanation of subset). To ask whether traits evolved under scenarios of Brownian motion (D = 0) or phylogenetic randomness (D = 1), observed values of D were compared to null distributions of D modeled under each scenario.

Evidence for correlated evolution was detected in two pairs of metabolite classes: 1) flavonoids and chalcones; and 2) *p*-alkenyl phenols and kavalactones/butenolides. For the first pair of traits, a model of contingency in which changes in chalcones depend on the state of flavonoids provided the best fit to the data (Table 2). In this model, when flavonoids are present, chalcone gains are almost two times more probable than chalcone losses; however, when flavonoids are absent, chalcone losses are much more probable than chalcone gains (Fig. 3). The alternative contingency model for this pair of traits (i.e., changes in flavonoids depend on the state of chalcone) was also a good fit to the data (Table 2). According to this model, when chalcones are present, flavonoid transitions are extremely probable, with flavonoid gains being approximately eight times more probable than flavonoid losses. Alternatively, when chalcones are absent, flavonoid losses are approximately five times more probable than flavonoid gains (Fig. 3). For the second pair of traits, *p*-alkenyl phenols and kavalactones/butenolides, the best fit model was one of interdependent correlated evolution in which changes in *p*-alkenyl phenol depend on the state of kavalactones/butenolides, and vice versa (Table 2). When kavalactones/butenolides are present, *p*-alkenyl phenol transitions are more probable than when they are absent, with the loss of *p*-alkenyl phenols being much more probable than the gain of *p*-alkenyl phenols under both scenarios. Alternatively, when *p*-alkenyl phenols are present, the loss of kavalactones/butenolides is extremely probable relative to the gain of kavalactones/butenolides, which is rarely observed. When *p*-alkenyl phenols are absent, kavalactones/butenolides are rarely gained or lost (Fig. 3).

**Table 2.**
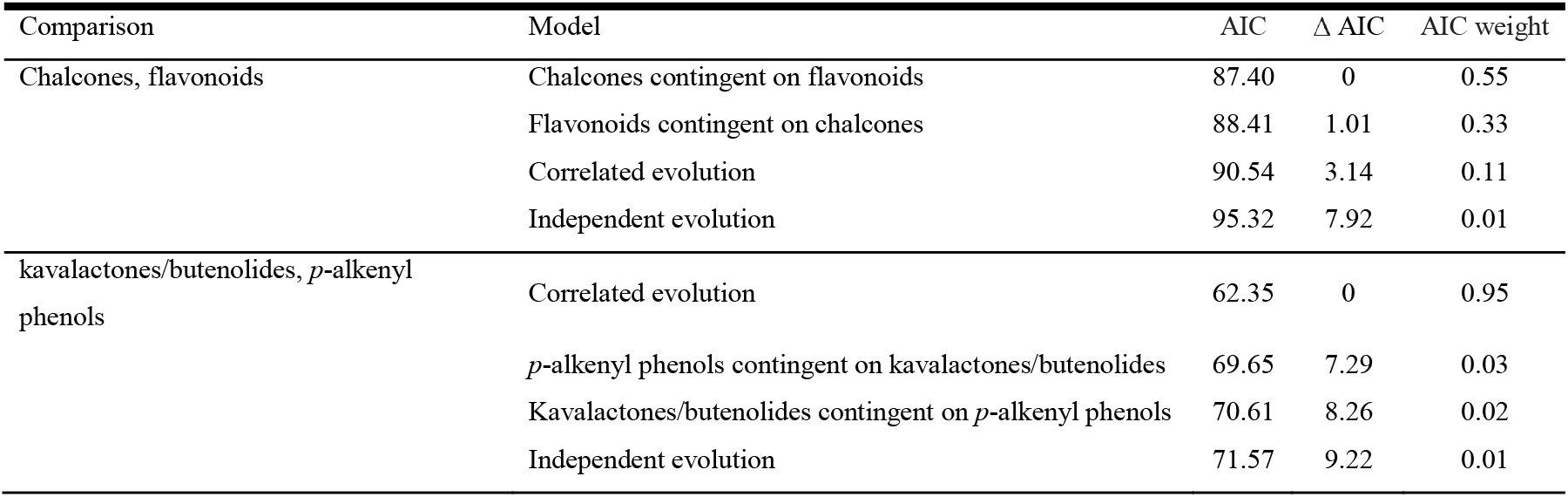
Correlated evolution was detected in two pairs of metabolite classes with Pagel’s (1994) method: 1) chalcones and flavonoids; and 2) kavalactones/butenolides and *p*-alkenyl phenols. A model comparison framework was employed to evaluate four potential models of trait evolution using AIC: correlated evolution (transition rate in one trait depends on state at another, and vice versa); contingent change (transition rate in one trait depends on state at another, but not the converse); and independent evolution.

| Comparison | Model | AIC | $\Delta$ AIC | AIC weight |
| --- | --- | --- | --- | --- |
| Chalcones, flavonoids | Chalcones contingent on flavonoids | 87.40 | 0 | 0.55 |
|  | Flavonoids contingent on chalcones | 88.41 | 1.01 | 0.33 |
|  | Correlated evolution | 90.54 | 3.14 | 0.11 |
|  | Independent evolution | 95.32 | 7.92 | 0.01 |
| kavalactones/butenolides, *p*-alkenyl phenols | Correlated evolution | 62.35 | 0 | 0.95 |
|  | *p*-alkenyl phenols contingent on kavalactones/butenolides | 69.65 | 7.29 | 0.03 |
|  | Kavalactones/butenolides contingent on *p*-alkenyl phenols | 70.61 | 8.26 | 0.02 |
|  | Independent evolution | 71.57 | 9.22 | 0.01 |

**Figure 3.**
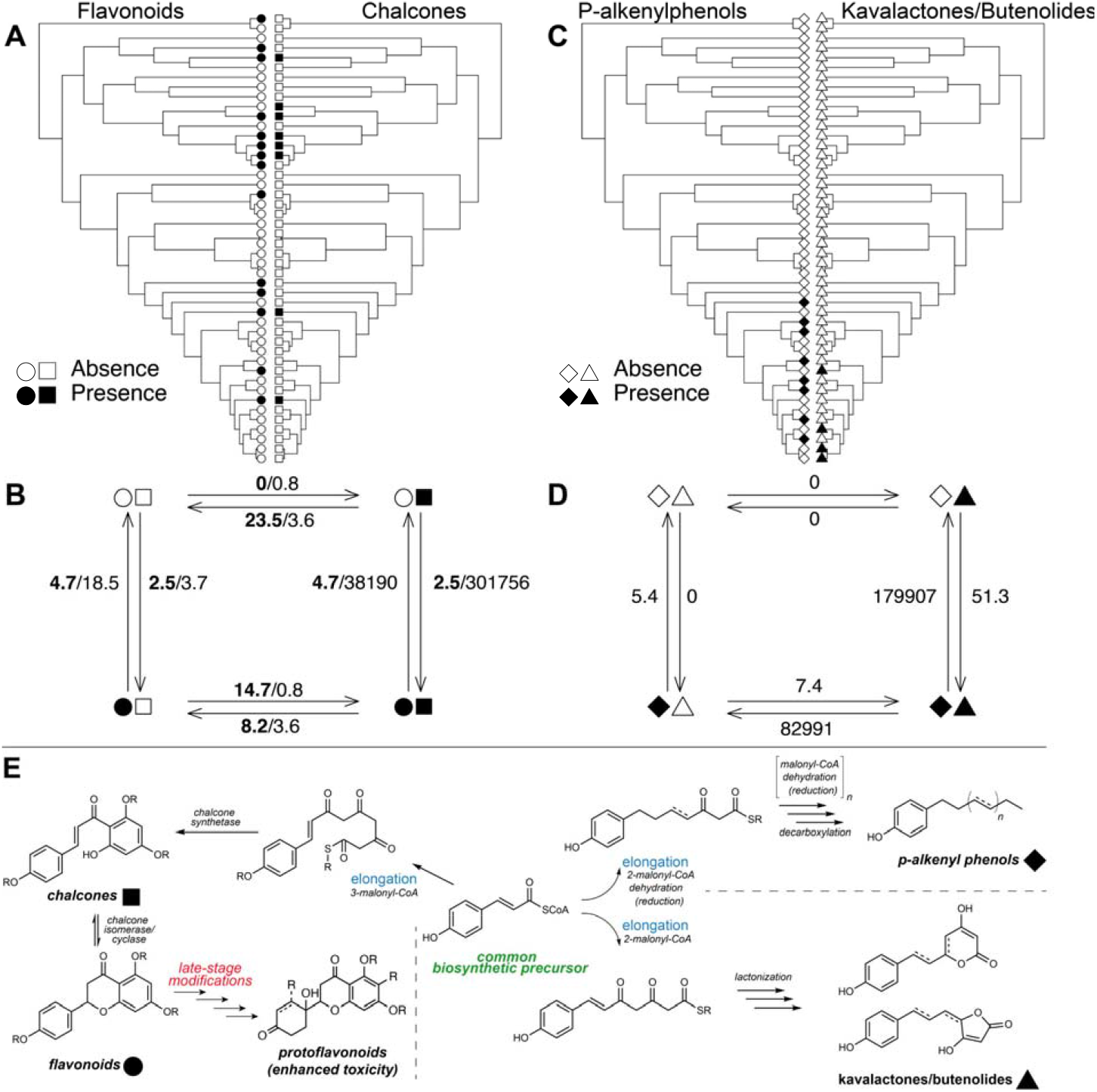
Evolutionary associations were detected in two pairs of traits according to Pagel’s (1994) test of correlated evolution: 1) flavonoids and chalcones and 2) *p*-alkenyl phenols and kavalactones/butenolides. Filled shapes indicate presences and unfilled shapes indicate absences of flavonoids (circles), chalcones (squares), *p*-alkenyl phenols (diamonds), and kavalactones/butenolides (triangles), respectively. The shapes used in the cophylogenetic plots (**A** and **C**) are repeated below (**B** and **D**) to depict four states comprising all combinations of presences and absences in the pair of traits. Arrows represent transition rates between states. **B.** As both models of contingent change provided good fits to the flavonoid and chalcone data, both sets of transition rates are displayed, with the first set of values (bolded) corresponding to the best supported model (chalcone evolution contingent on flavonoid state) and the second set of values corresponding to the alternative contingency model (flavonoid evolution contingent on chalcone state). **D.** The best fit model to the *p*-alkenyl phenol and kavalactone/butenolide data was one of dependent evolution, where *p*-alkenyl phenol evolution is dependent on the state at the kavalactone/butanolide trait, and vice versa. Panel **E** illustrates the enzymatic processes and branch points along biosynthetic pathways that give rise to the four classes of metabolites. Chalcones are immediate biosynthetic precursors of flavonoids, where the inherent reactivity of the chalcone moiety permits cyclization to the flavonoid scaffold. Subtle structural changes to the flavonoid scaffold caused by late-stage oxidation can produce protoflavonoids, a rare class of metabolite with potent cytotoxic activity. In contrast, the pathways of *p-*alkenyl phenols and kavalactones diverge much earlier and embark on distinct chain elongation pathways which lead to long-chain lipophilic substituent characteristic of the *p*-alkenyl phenols in one case, and lactones (kavalactones and butenolides) in the other case.

### Phylogenetic signal in high-dimensional metabolomic data

While broad metabolite classes uniformly exhibited at least moderate levels of phylogenetic signal, evidence for phylogenetic signal in multivariate analyses of the crude ^1^H NMR data was mixed. PCs 1 & 2 and 3 & 4 explained 47.89% and 17.16% of variance in the ^1^H NMR data, respectively, but showed little clustering by clade (Fig. 4a). Permutational multivariate analyses of variance were not significant for combinations of neither PC 1 & 2 (*P* = 0.635) nor 3 & 4 (*P* = 0.445), suggesting that different clades do not form distinct clusters in chemospace based on their ^1^H NMR spectra.

**Figure 4.**
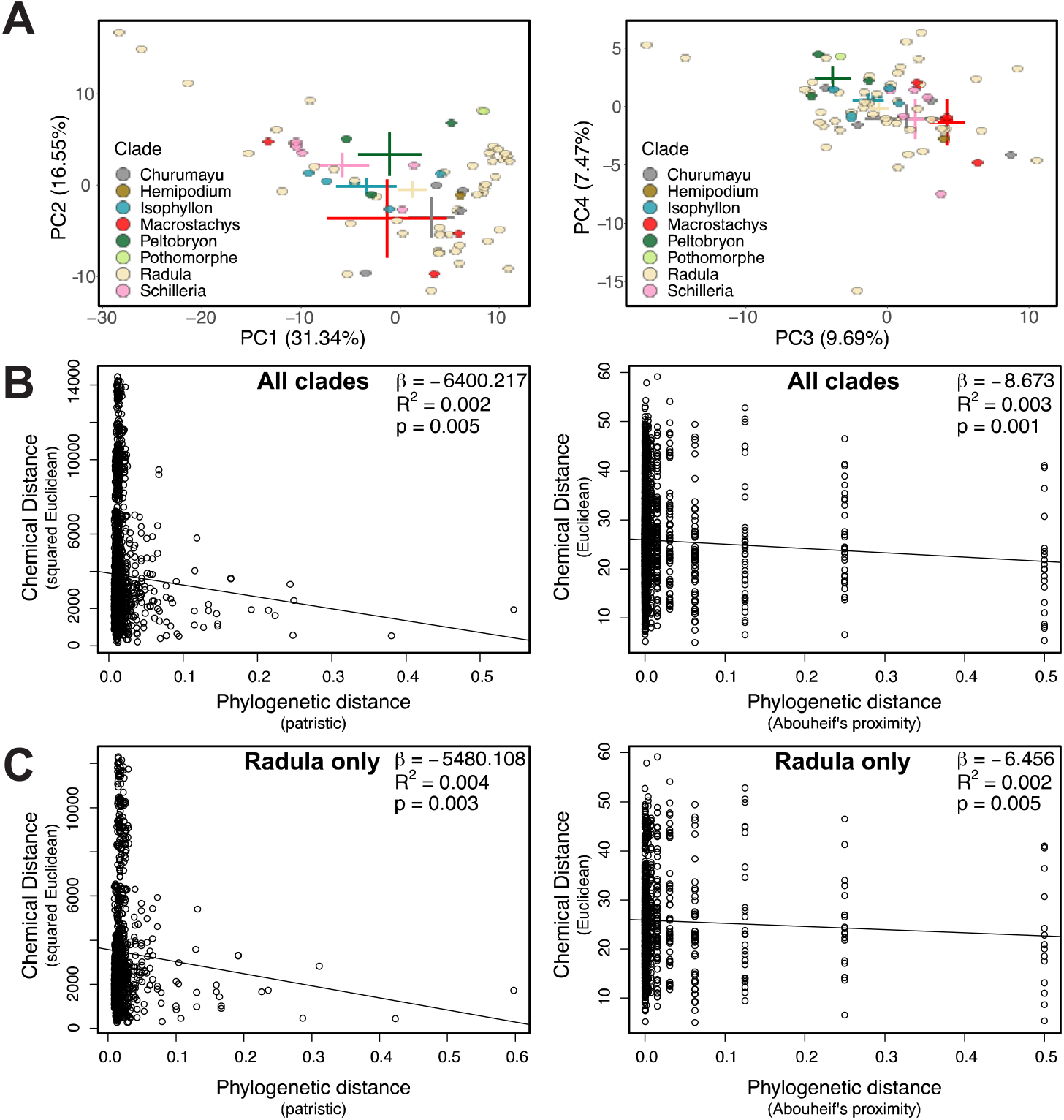
**A.** Chemospace of all 71 species constructed with the crude ^1^H NMR data across 277 peaks. Point shapes and colors are formatted according to clade designation as portrayed in the phylogenetic tree in Figure 1. MRM analyses recovered significant negative relationships between phylogenetic and chemical distances calculated among samples from all clades (**B**), or from the Radula clade only (**C**); however, the proportion of variance explained was low for all tests.

According to the MRM models, both patristic distance and Abouheifs proximity significantly predict a small proportion of variation in chemical distance calculated among *Piper* samples from all clades (patristic: β = −6400.217, *R*^2^ = 0.002, *P* = 0.005; Abouheif: β = −8.673, *R*^2^ = 0.003, *P* = 0.001) and among Radula samples only (patristic: β = −5480.108, *R*^2^ = 0.004, *P* = 0.003; Abouheif: β = −6.456, *R*^2^ = 0.002, *P* = 0.005) (Fig. 4bc). Though explained variance is small, the slope coefficients for these significant relationships are negative, indicating that decreasing phylogenetic distance is associated with increasing chemical distance.

Analyses with the generalized *K* statistic (*K*_mult_; Adams, 2014) indicated lower levels of phylogenetic signal in the metabolomic data than expected under a Brownian motion model of evolution for *Piper* generally (*K*_mult_ = 0.1606, *P* = 0.001) and for Radula specifically (*K*_mult_ = 0.1803, *P* = 0.001). Still, the observed *K*_mult_ was higher than all *K*_mult_ values obtained with permutations of the ^1^H NMR dataset (Fig. S1). Additionally, few *K*_mult_ tests of the permuted data yielded significant *P*-values (4.4% of permutations), indicating that the estimate we observed, though subtle and lower than Brownian motion expectations, was real and not a statistical artifact of zero-inflation in the data.

## Discussion

*Piper* is a hyper-diverse lineage in which phytochemical variation has influenced evolutionary and ecological processes and shaped complex tropical communities (e.g., Salazar *et al*., 2016; Richards *et al*., 2018). However, there have been limitations in both the degree of phylogenetic resolution and the understanding of phytochemical variation in this group. Phylogenies inferred here with ddRADseq data substantially improved resolution and support compared to past studies of *Piper*, which were limited by interspecific variation in small numbers of Sanger-sequenced loci (Jaramillo *et al*., 2008; Smith *et al*., 2008; Martínez *et al*., 2015). Although the data set did not include members from all previously recognized groups, analyses resolved eight monophyletic Neotropical *Piper* clades, six of which have been inferred in previous analyses of the genus based on chloroplast psbJ-petA and ITS (Jaramillo *et al*., 2008; Martínez *et al*., 2015). Two of the eight clades, Churumayu and Isophyllon, had been previously nested within Radula (Jaramillo *et al*., 2008); however, our results suggest that they are independent monophyletic lineages (Fig. 1). Despite low support for several deep divergences, the phylogeny inferred here had strong resolution and support for recent relationships, including within Radula (Fig. 1), consistent with other recent reduced representation sequencing studies that have generated high quality phylogenies at shallow time scales (Massati *et al*., 2016; Eaton *et al*., 2017; Lecaudey *et al*., 2018; Paetzold *et al*., 2019). However, a potential limitation of such sequencing designs may include the recovery of fewer loci shared by more distantly related samples due to allelic dropout (Cariou *et al*., 2013; Cooke *et al*., 2016). It is possible that allelic dropout, potentially acerbated by strict filtering of missing data, led to weak support values for deep splits in the phylogeny, many of which occurred early in the history of the Neotropical *Piper* lineage (Martínez *et al*., 2015). Nonetheless, the resulting subset of data (641 loci; 9,113 SNPs) was sufficient for inferring a largely resolved phylogeny, highlighting the potential promise of reduced representation sequencing for resolving evolutionary histories even in groups spanning moderately deep divergence.

Comparative studies have taken diverse approaches to analyzing metabolomic data, each providing a unique perspective on the evolution of specialized metabolites (e.g., Salazar *et al*., 2018; Sedio *et al*., 2018, 2019; Ernst *et al*., 2019; Kang *et al*., 2019). Here, we first characterized the presence/absence of 35 metabolite classes commonly used to categorize plant secondary compounds that are hierarchically nested into three levels of structural resolution. Specific categories at the lowest level of the hierarchy, representing specialized structural motifs or specific molecules, were rare across species and precluded tests of phylogenetic signal at our level of taxonomic sampling (Fig. 2). Despite not being able to test for phylogenetic signal, clustering is evident for more specific categories, such as crassinervic acid and prenylated flavonoids, which are only present in small subclades but include particularly effective defenses (Dyer & Palmer, 2004; Salehi *et al*., 2019). Alternatively, broader metabolite classes at intermediate and high positions in the hierarchy that are directly tied to fundamental secondary metabolite biosynthetic pathways were more abundant across species and exhibited moderate-high levels of phylogenetic signal across Radula (Table 1, Fig. 2). This pattern may be expected if initial biosynthetic steps are conserved over longer evolutionary scales, permitting the abundance of broad chemical classes, yet later stage modifications of these core structures are more evolutionarily labile, causing structural similarity to be low even among related species. Flavonoids are a good example of this pattern, with pathways that form the flavonoid scaffold being very conserved, as they are catalyzed by modified enzymes from ubiquitous metabolic pathways, but then subsequent biosynthetic steps (e.g., those catalyzed by p450 enzymes) modify these scaffolds (Yonekura-Sakakibara *et al*., 2019), yielding unique molecules towards the tips of evolutionary trees (Fig. 3E). For example, late-stage modification of common flavonoid scaffolds can result in the production of non-aromatic protoflavanoids. These compounds rarely occur across the plant kingdom and have only recently been found in one species of *Piper* (Freitas *et al*., 2014). Importantly, this subtle structural modification that leaves most of the flavonoid scaffold intact has been demonstrated to dramatically enhance the cytotoxic properties compared to that of the parent flavonoid (Hunyadi *et al*., 2014; Latif *et al*., 2020).

One key prediction from the escape and radiate hypothesis is that adaptive defensive traits should be phylogenetically conserved within the lineage they evolved, but this prediction has mostly been evaluated with broad classes of secondary metabolites at high taxonomic scales (e.g., Ehrlich & Raven, 1964; Moreira *et al*., 2018; Yonekura-Sakakibara *et al*., 2019; Zhang *et al*., 2020) rather than specific compounds in recent diversifications (e.g., Agrawal *et al*., 2009; Salazar *et al*., 2018; Allevato *et al*., 2019). A growing number of studies conducted at shallower evolutionary scales suggest chemical traits may be evolutionarily labile and highlight the need for determining the level at which chemical defense is conserved, and which compound classes are more likely to exhibit phylogenetic signal and evolutionary correlations (Kursar *et al*., 2009; Sedio, 2013; Johnson *et al*., 2014; Salazar *et al*., 2016; Maldonado *et al*., 2017; Moreira *et al*., 2018). Further, an understanding of the phylogenetic scale of chemical trait conservation will enable insights into the drivers of herbivorous insect radiations, as the nature of codiversification in many of these lineages is likely structured by complex associations between geology, geography, chemical defense, and biotic interactions (Endara *et al*., 2017; Jahner *et al*., 2017). Our results are generally consistent with the predictions of signal (and conservatism) for broad classes of compounds, as well as the lack of signal for specific structures captured by ^1^H NMR data.

The ^1^H NMR data address a different set of hypotheses than data from categorization of individual molecules – peaks represent resonances associated with particular molecular structures rather than individual compounds, and the chemical shift (frequency), shape, and abundance of these resonances are extremely sensitive to subtle structural changes. ^1^H NMR spectroscopy easily detects a great range and subtle differences in compositional and structural complexity, including increasing size, asymmetry and oxidation states, that might be predicted to evolve in response to divergent selection across plant populations responding to different suites of enemies (Dyer *et al*., 2018). Low levels of phylogenetic signal in the ^1^H NMR data and evidence for phylogenetic overdispersion (Fig. 4) is also likely due to the fact that many molecular features of small defensive molecules have potentially evolved in a convergent manner across *Piper*, such as the kavalactones, *p*-alkenyl phenols, piplartine, oxidized prenylated benzoic acids, chromanes, anuramide peptides, and phenethyl amides.

There are numerous limitations that could affect estimates of phylogenetic signal in comparative studies (reviewed by Kamilar & Cooper, 2013) that are relevant to the analyses presented here. First, incomplete taxon sampling and unresolved tree structure can substantially influence tests of phylogenetic signal and likely influenced our results to some degree. However, we made great effort to sample species from across the entire known phylogeny of Radula to reduce sampling bias, and more comprehensive genomic sampling produced enhanced phylogenetic resolution of the Radula clade, where we focused the majority of phylogenetic comparative methods. In addition, we were unable to quantify the measurement error associated with the chemical traits within species (e.g., Johnson *et al*., 2014), which can decrease the statistical power for detecting phylogenetic signal (Blomberg *et al*., 2003; Ives *et al*., 2007; Hardy & Pavoine, 2012). It is also possible that environmental effects on our chemical traits could bias estimates of phylogenetic signal and correlations (Ives *et al*., 2007).

The causes of correlated evolution, including linkage, epistasis, and selection, are difficult to detect without careful approaches in quantitative genetics and population genomics. Nevertheless, one advantage of examining the presence/absence of multiple classes of defensive compounds in a phylogenetic context is that it is possible to test for expected patterns of correlated evolution due to shared metabolic pathways (e.g., flavonoids and cardenolides; Agrawal *et al*., 2009) or due to adaptive advantages of specific mixtures. Recent studies detecting evolutionary associations among chemical traits (Johnson *et al*., 2014; Kariñho-Betancourt *et al*., 2015; Boachon *et al*., 2018) have posited that the branching structure of metabolic pathways could potentially drive this pattern. If metabolite classes share a common precursor, one might expect evolutionary tradeoffs and negative covariation. Alternatively, if metabolite classes lie along the same metabolic pathway, an increase in one class may be concomitant with increases in another (or vice versa) causing positive covariation among the classes. There are also numerous empirical examples supporting the hypotheses that positive correlations may be driven by functional redundancy (Jones & Firn, 1991; Romeo *et al*., 2013) or selection for synergistic effects on herbivores (Dyer *et al*., 2003; Richards *et al*., 2010) rather than the structural constraints of metabolism. Suites of covarying defensive traits, or defense syndromes, have been detected in several plant genera (Becerra *et al*., 2001; Agrawal & Fishbein, 2006; Endara *et al*., 2017) and plant communities (Kursar & Coley, 2003), and have been predominantly used to describe covariation among mechanical and chemical defenses. It is interesting to note the correlated evolution of the flavones/chalcones and the *p*-alkenyl phenols/kavalactones could be due to metabolic constraints, as well as possible adaptations via synergistic (e.g., kavalactones in *P. methysticum)* or other mixture-associated defensive attributes (reviewed in Dyer *et al*., 2018). Flavonoids and chalcones are directly linked biosynthetically, such that the inherent reactivity of the chalcone moiety permits the enzymatic processes that result in cyclization to the flavonoid scaffold (Fig. 3E). This strong biosynthetic tie predicts the presence of one would depend on the other, and indeed our structural analysis found many cases where both metabolite classes co-occurred in the same sample. Revealing the relationship between the kavalactones and *p*-alkenyl phenols is more tenuous because both classes are less prevalent across our samples. Kavalactones and *p*-alkenyl phenols are dramatically different compounds that diverge at a much earlier branch point from a common cinnamic/coumaric acid precursor. Whereas one polyacetate chain extension pathway leads to the long-chain lipophilic substituent, characteristic of the *p*-alkenyl phenols, the other chain extension pathway conserves oxidation states through the chain extension process to produce the lactones (kavalactones or butenolides) through cyclization reactions (Fig. 3E). The overall outcome is different than the chalcone-flavonoid relationship; in this case, two dramatically different compounds are produced by divergence from a common early-stage biosynthetic precursor in contrast to the immediate biosynthetic precursor relationship between chalcones and flavonoids. Broader sampling across *Piper* and Radula will be necessary to confirm this unexpected relationship between kavalactones and *p*-alkenyl phenols.

## Conclusion

Here we sought to advance understanding of phylogenetic relationships within *Piper* while simultaneously investigating the mode and manner of phytochemical evolution in this group. In addition to generating a well-resolved phylogeny, our results support theoretical expectations that broad classes of compounds display higher degrees of phylogenetic conservatism than the more evolutionarily labile molecular features revealed by ^1^H NMR data. In addition, trait associations observed in Radula can be used to pose functional hypotheses about genetic constraints or biases on phytochemical evolution and how these factors structure plant-animal interactions. Such investigations are one of the emerging frontiers in terrestrial ecology, and we hope that our study provides one example of how collaborative and multi-disciplinary research can progress in this area.

## Supplementary material

**Table S1.** Sampling information for all individuals.

**Fig. S1.** Results of the multivariate K test on 1000 permutations of all chemical regions.

## Acknowledgements

This research was funded by the National Science Foundation (DEB-1145609 and DEB-1442103) to CJ, LAD, LAR, MLF, TLP, and AMS, by the National Science Foundation Graduate Research Award (Award No. 1650114) to KAU, and by FAPESP (Award No 2014/50316-7) to MJK. Fellowship support for KAU, KMO, and CSP and funding for chemical instrumentation and analysis was provided by the Hitchcock Center for Chemical Ecology at the University of Nevada, Reno. We thank Jennifer L. McCracken for her assistance with the collection of GC-MS data for the categorical chemical characterization, and we thank Chris Feldman, Beth Leger, and Steve Vander Wall for their guidance during the earliest stages of this project.

## Author contributions

MLF, LAD, AMS, CSJ, LAR, and TLP developed the original idea for the research and secured funding. EJT, MJK, and LFY collected specimens. EJT extracted DNA from plant specimens. KAU and TLP generated genotyping-by-sequencing libraries. KAU and JPJ analyzed the genetic data. KMO and LAR performed chemical extractions and analyses. CSJ, CSP, and CDD executed chemical annotation and structure determination. KAU and JPJ wrote the first draft of the manuscript, and all authors contributed to subsequent revisions.

## References

Abouheif E. 1999. A method for testing the assumption of phylogenetic independence in comparative data. Evolutionary Ecology Research 1: 895–909.

Adams DC. 2014. A generalized *K* statistic for estimating phylogenetic signal from shape and other high-dimensional multivariate data. Systematic Biology 63: 685–697.

Adams DC, Otárola Castillo E. 2013. geomorph: an R package for the collection and analysis of geometric morphometric shape data. Methods in Ecology and Evolution 4: 393–399.

Agrawal AA. 2007. Macroevolution of plant defense strategies. Trends in Ecology & Evolution 22:103–109.

Agrawal AA, Fishbein M. 2006. Plant defense syndromes. Ecology 87: S132–S149.

Agrawal AA, Salminen JP, Fishbein M. 2009. Phylogenetic trends in phenolic metabolism of milkweeds *(Asclepias):* evidence for escalation. Evolution: International Journal of Organic Evolution 63: 663–673.

Allevato DM, Groppo M, Kiyota E, Mazzafera P, Nixon KC. 2019. Evolution of phytochemical diversity in *Pilocarpus* (Rutaceae). Phytochemistry 163:132–146.

Anderson MJ. 2001. A new method for non parametric multivariate analysis of variance. Austral Ecology 26: 32–46.

Andrews KR, Good JM, Miller MR, Luikart G, Hohenlohe PA. 2016. Harnessing the power of RADseq for ecological and evolutionary genomics. Nature Reviews Genetics 17: 81–92.

Asmarayani R. 2018. Phylogenetic relationships in Malesian–Pacific *Piper* (Piperaceae) and their implications for systematics. Taxon 67: 693–724.

Bagley JC, Uribe-Convers S, Carlsen MM, Muchhala N. 2020. Utility of targeted sequence capture for phylogenomics in rapid, recent angiosperm radiations: Neotropical *Burmeistera* bellflowers as a case study. Molecular Phylogenetics and Evolution 152: 106769.

Becerra JX. 1997. Insects on plants: macroevolutionary chemical trends in host use. Science 276: 253–256.

Becerra JX, Venable D, Evans P, Bowers W. 2001. Interactions between chemical and mechanical defenses in the plant genus *Bursera* and their implications for herbivores. American Zoologist 41: 865–876.

Berenbaum M. 1978. Toxicity of a furanocoumarin to armyworms: a case of biosynthetic escape from insect herbivores. Science 201: 532–534.

Blomberg SP, Garland T, Ives AR. 2003. Testing for phylogenetic signal in comparative data: behavioral traits are more labile. Evolution 57: 717–745.

Boachon B, Buell CR, Crisovan E, Dudareva N, Garcia N, Godden G, Henry L, Kamileen MO, Kates HR, Kilgore MB et al. 2018. Phylogenomic mining of the mints reveals multiple mechanisms contributing to the evolution of chemical diversity in Lamiaceae. Molecular Plant 1: 1084–1096.

Bowers MD. 1983. The role of iridoid glycosides in host-plant specificity of checkerspot butterflies. Journal of Chemical Ecology 9: 475–493.

Bowers MD. 1984. Iridoid glycosides and host-plant specificity in larvae of the buckeye butterfly, *Junonia coenia* (Nymphalidae). Journal of Chemical Ecology 10: 1567–1577.

Callejas-Posada R. 2020. Piperaceae. In: Davidse G, Ulloa Ulloa C, Hernández Macías HM, Knapp S, eds. Flora Mesoamericana, vol. 2, pt. 2 (pp. i–xix; 1–590), St. Louis, MO: Missouri Botanical Garden Press.

Cardini A, Elton S. 2008. Does the skull carry a phylogenetic signal? Evolution and modularity in the guenons. Biological Journal of the Linnean Society 93: 813–834.

Cariou M, Duret L, Charlat S. 2013. Is RAD□seq suitable for phylogenetic inference? An in silico assessment and optimization. Ecology and Evolution. 3:846–852.

Carter KA, Liston A, Bassil NV, Alice LA, Bushakra JM, Sutherland BL, Mockler TC, Bryant DW, Hummer KE. 2019. Target capture sequencing unravels *Rubus* evolution. Frontiers in Plant Science 10: 1615.

Caseys C, Stritt C, Glauser G, Blanchard T, Lexer C. 2015. Effects of hybridization and evolutionary constraints on secondary metabolites: the genetic architecture of phenylpropanoids in European *Populus* species. PloS one 10: e0128200.

Chen F, Tholl D, Bohlmann J, Pichersky E. 2011. The family of terpene synthases in plants: a midsize family of genes for specialized metabolism that is highly diversified throughout the kingdom. The Plant Journal 66: 212–229.

Clavel J, Escarguel G, Merceron G. 2015. mvmorph: an R package for fitting multivariate evolutionary models to morphometric data. Methods in Ecology and Evolution 6: 1311–1319.

Colby S, Alonso W, Katahira E, McGarvey D, Croteau R. 1993. 4s-limonene synthase from the oil glands of spearmint *(Mentha spicata).* cDNA isolation, characterization, and bacterial expression of the catalytically active monoterpene cyclase. Journal of Biological Chemistry 268:23016–23024.

Cooke TF, Yee MC, Muzzio M, Sockell A, Bell R, Cornejo OE, Kelley JL, Bailliet G, Bravi CM, Bustamante CD, Kenny EE. 2016. GBStools: a statistical method for estimating allelic dropout in reduced representation sequencing data. PLoS Genetics 12: e1005631.

Du ZY, Harris AJ, Xiang QYJ. 2020. Phylogenomics, co-evolution of ecological niche and morphology, and historical biogeography of buckeyes, horsechestnuts, and their relatives (Hippocastaneae, Sapindaceae) and the value of RAD-seq for deep evolutionary inferences back to the Late Cretaceous. Molecular Phylogenetics and Evolution 145: 106726.

Dyer LA, Dodson CD, Stireman JO, Tobler MA, Smilanich AM, Fincher RM, Letourneau DK. 2003. Synergistic effects of three *Piper* amides on generalist and specialist herbivores. Journal of Chemical Ecology 29: 2499–2514.

Dyer LA, Palmer AD. 2004. Piper: a model genus for studies of phytochemistry, ecology, and evolution. New York, NY: Kluwer Academic/Plenum Publishers.

Dyer LA, Philbin CS, Ochsenrider KM, Richards LA, Massad TJ, Smilanich AM, Forister ML, Parchman TL, Galland LM, Hurtado PJ, et al. 2018. Modern approaches to study plant–insect interactions in chemical ecology. Nature Reviews Chemistry 2: 50–64.

Dyer LA, Richards J, Dodson CD. 2004. Isolation, synthesis, and evolutionary ecology of *Piper* amides. In: Dyer LA, Palmer AD, eds. Piper: A model genus for studies of phytochemistry, ecology, and evolution. City, State: Springer, 117–139.

Easson CG, Thacker RW. 2014. Phylogenetic signal in the community structure of host-specific microbiomes of tropical marine sponges. Frontiers in Microbiology 5: 532.

Eaton DA. 2014. PyRAD: assembly of *de novo* RADseq loci for phylogenetic analyses. Bioinformatics 30: 1844–1849.

Eaton DA, Ree RH. 2013. Inferring phylogeny and introgression using RADseq data: an example from flowering plants *(Pedicularis:* Orobanchaceae). Systematic Biology 62: 689–706.

Eaton DA, Spriggs EL, Park B, Donoghue MJ. 2017. Misconceptions on missing data in RADseq phylogenetics with a deep-scale example from flowering plants. Systematic Biology 66: 399–412.

Edgar RC. 2004. MUSCLE: multiple sequence alignment with high accuracy and high throughput. Nucleic Acids Research 32: 1792–1797.

Ehrlich PR, Raven PH. 1964. Butterflies and plants: a study in coevolution. Evolution 18: 586–608.

Endara MJ, Coley PD, Ghabash G, Nicholls JA, Dexter KG, Donoso DA, Stone GN, Pennington RT, Kursar TA. 2017. Coevolutionary arms race versus host defense chase in a tropical herbivore-plant system. Proceedings of the National Academy of Sciences USA 114: E7499–E7505.

Ernst M, Nothias LF, van der Hooft JJJ, Silva RR, Saslis-Lagoudakis CH, Grace OM, Martinez-Swatson K, Hassemer G, Funez LA, Simonsen HT, et al. 2019. Assessing specialized metabolite diversity in the cosmopolitan plant genus *Euphorbia* L. Frontiers in Plant Science 10: 846.

Felsenstein J. 1985. Phylogenies and the comparative method. The American Naturalist 125: 1–15.

Fernández-Mazuecos M, Mellers G, Vigalondo B, Sáez L, Vargas P, Glover BJ. 2017. Resolving recent plant radiations: power and robustness of genotyping-by-sequencing. Systematic Biology 67: 250–268.

Fine PVA, Miller ZJ, Mesones I, Irazuzta S, Appel HM, Stevens MHH, Sääksjärvi I, Schultz JC, Coley PD. 2006. The growth–defense trade-off and habitat specialization by plants in Amazonian forests. Ecology 87: S150–S162.

Freitas GC, Batista Jr JM, Franchi Jr GC, Nowill AE, Yamaguchi LF, Vilcachagua JD, Favaro DC, Furlan M, Guimarães EF, Jeffrey CS, Kato MJ. 2014. Cytotoxic non-aromatic B-ring flavanones from *Piper carniconnectivum* C. DC. Phytochemistry 97: 81–87.

Fritz SA, Purvis A. 2010. Selectivity in mammalian extinction risk and threat types: a new measure of phylogenetic signal strength in binary traits. Conservation Biology 24:1042–1051.

Fukushima A, Kusano M, Redestig H, Arita M, Saito K. 2011. Metabolomic correlation-network modules in *Arabidopsis* based on a graph-clustering approach. BMC Systems Biology 5: 1.

Gentry AH. 1993. Four neotropical rainforests. New Haven, CT: Yale University Press.

Glassmire AE, Jeffrey CS, Forister ML, Parchman TL, Nice CC, Jahner JP, Wilson JS, Walla TR, Richards LA, Smilanich AM, Leonard MD. 2016. Intraspecific phytochemical variation shapes community and population structure for specialist caterpillars. New Phytologist 212: 208–219.

Goslee SC, Urban DL. 2007. The ecodist package for dissimilarity-based analysis of ecological data. Journal of Statistical Software 22: 1–19.

Griffin WJ, Lin GD. 2000. Chemotaxonomy and geographical distribution of tropane alkaloids. Phytochemistry 53: 623–637.

Hamon P, Grover CE, Davis AP, Rakotomalala J-J, Raharimalala NE, Albert VA, Sreenath HL, Stoffelen P, Mitchell SE, Couturon E, et al. 2017. Genotyping-by-sequencing provides the first well-resolved phylogeny for coffee *(Coffea)* and insights into the evolution of caffeine content in its species. Molecular Phylogenetics and Evolution 109: 351–361.

Hardy OJ, Pavoine S. 2012. Assessing phylogenetic signal with measurement error: a comparison of Mantel tests, Blomberg et al.’s *K*, and phylogenetic distograms. Evolution: International Journal of Organic Evolution 66: 2614–2621.

Harmon LJ, Glor RE. 2010. Poor statistical performance of the Mantel test in phylogenetic comparative analyses. Evolution: International Journal of Organic Evolution 64: 2173–2178.

Herrera S, Shank TM. 2016. RAD sequencing enables unprecedented phylogenetic resolution and objective species delimitation in recalcitrant divergent taxa. Molecular Phylogenetics and Evolution 100: 70–79.

Hipp AL, Manos PS, Hahn M, Avishai M, Bodénès C, Cavender-Bares J, Crowl AA, Deng M, Denk T, Fitz-Gibbon S, Gailing O. 2020. Genomic landscape of the global oak phylogeny. New Phytologist 226: 1198–1212.

Höhna S, Landis MJ, Heath TA, Boussau B, Lartillot N, Moore BR, Huelsenbeck JP, Ronquist F. 2016. RevBayes: Bayesian phylogenetic inference using graphical models and an interactive model-specification language. Systematic Biology 65: 726–736.

Hunyadi A, Martins A, Danko B, Chang FR, Wu YC. 2014. Protoflavones: A class of unusual flavonoids as promising novel anticancer agents. Phytochemistry Reviews 13: 69–77.

Ives AR, Midford PE, Garland Jr T. 2007. Within-species variation and measurement error in phylogenetic comparative methods. Systematic Biology 56: 252–270.

Jahner JP, Forister ML, Parchman TL, Smilanich AM, Miller JS, Wilson JS, Walla TR, Tepe EJ, Richards LA, Quijano-Abril MA, et al. 2017. Host conservatism, geography, and elevation in the evolution of a Neotropical moth radiation. Evolution 71: 2885–2900.

Janz N. 2011. Ehrlich and Raven revisited: mechanisms underlying codiversification of plants and enemies. Annual Review of Ecology, Evolution, and Systematics 42: 71–89.

Jaramillo MA, Callejas R, Davidson C, Smith JF, Stevens AC, Tepe EJ. 2008. A phylogeny of the tropical genus *Piper* using ITS and the chloroplast intron psbJ–petA. Systematic Botany 33: 647–660.

Johnson MT, Agrawal AA, Maron JL, Salminen JP. 2009. Heritability, covariation and natural selection on 24 traits of common evening primrose *(Oenothera biennis)* from a field experiment. Journal of Evolutionary Biology 22: 1295–1307.

Johnson MT, Ives AR, Ahern J, Salminen JP. 2014. Macroevolution of plant defenses against herbivores in the evening primroses. New Phytologist 203: 267–279.

Jombart T, Devillard S, Balloux F. 2010. Discriminant analysis of principal components: a new method for the analysis of genetically structured populations. BMC Genetics 11: 94.

Jones CG, Firn RD. 1991. On the evolution of plant secondary chemical diversity. Philosophical Transactions of the Royal Society of London. Series B: Biological Sciences 333: 273–280.

Jost L. 2006. Entropy and diversity. Oikos 113: 363–375.

Kamilar JM, Cooper N. 2013. Phylogenetic signal in primate behaviour, ecology and life history. Philosophical Transactions of the Royal Society B: Biological Sciences 368: 20120341.

Kang KB, Ernst M, van der Hooft JJJ, da Silva RR, Park J, Medema MH, Sung SH, Dorrestein PC. 2019. Comprehensive mass spectrometry-guided phenotyping of plant specialized metabolites reveals metabolic diversity in the cosmopolitan plant family Rhamnaceae. The Plant Journal 98:1134–1144.

Kariñho-Betancourt E, Agrawal AA, Halitschke R, Núñez-Farfán J. 2015. Phylogenetic correlations among chemical and physical plant defenses change with ontogeny. New Phytologist 206: 796–806.

Kato MJ, Furlan M. 2007. Chemistry and evolution of the Piperaceae. Pure and Applied Chemistry 79: 529–538.

Klingenberg CP, Gidaszewski NA. 2010. Testing and quantifying phylogenetic signals and homoplasy in morphometric data. Systematic Biology 59: 245–261.

Kursar TA, Coley PD. 2003. Convergence in defense syndromes of young leaves in tropical rainforests. Biochemical Systematics and Ecology 31: 929–949.

Kursar TA, Dexter KG, Lokvam J, Pennington RT, Richardson JE, Weber MG, Murakami ET, Drake C, McGregor R, Coley PD. 2009. The evolution of antiherbivore defenses and their contribution to species coexistence in the tropical tree genus *Inga*. Proceedings of the National Academy of Sciences 106: 18073–18078.

Langmead B, Salzberg SL. 2012. Fast gapped-read alignment with Bowtie 2. Nature methods 9: 357–359.

Latif AD, Jernei T, Podolski-Renić A, Kuo CY, Vágvölgyi M, Girst G, Zupkó I, Develi S, Ulukaya E, Wang HC, Pešić M, Csámpai A and Hunyadi A. 2020. Protoflavone-chalcone hybrids exhibit enhanced antitumor action through modulating redox balance, depolarizing the mitochondrial membrane, and inhibiting ATR-dependent signaling. Antioxidants 9: 1–18.

Leaché AD, Oaks JR. 2017. The utility of single nucleotide polymorphism (SNP) data in phylogenetics. Annual Review of Ecology, Evolution, and Systematics 48: 69–84.

Lecaudey LA, Schliewen UK, Osinov AG, Taylor EB, Bernatchez L, Weiss SJ. 2018. Inferring phylogenetic structure, hybridization and divergence times within Salmoninae (Teleostei: Salmonidae) using RAD-sequencing. Molecular Phylogenetics and Evolution 124: 82–99.

Léveillé-Bourret É, Chen BH, Garon-Labrecque MÉ, Ford BA, Starr JR. 2020. RAD sequencing resolves the phylogeny, taxonomy and biogeography of Trichophoreae despite a recent rapid radiation (Cyperaceae). Molecular Phylogenetics and Evolution 145: 106727.

Malcolm SB. 1994. Milkweeds, monarch butterflies and the ecological significance of cardenolides. Chemoecology 5: 101–117.

Maldonado C, Barnes CJ, Cornett C, Holmfred E, Hansen SH, Persson C, Antonelli A, Rønsted N. 2017. Phylogeny predicts the quantity of antimalarial alkaloids within the iconic yellow Cinchona bark (Rubiaceae: *Cinchona calisaya)*. Frontiers in Plant Science 8: 391.

Maron JL, Agrawal AA, Schemske DW. 2019. Plant-herbivore coevolution and plant speciation. Ecology 100: e02704.

Martínez C, Carvalho MR, Madriñán S, Jaramillo CA. 2015. A late Cretaceous *Piper* (Piperaceae) from Colombia and diversification patterns for the genus. American Journal of Botany 102: 273–289.

Massatti R, Reznicek AA, Knowles LL. 2016. Utilizing RADseq data for phylogenetic analysis of challenging taxonomic groups: A case study in *Carex* sect. *Racemosae*. American Journal of Botany 103: 337–347.

Mithöfer A, Boland W. 2012. Plant defense against herbivores: chemical aspects. Annual Review of Plant Biology 63: 431–450.

Molina-Henao YF, Guerrero-Chacón AL, Jaramillo MA. 2016. Ecological and geographic dimensions of diversification in *Piper* subgenus *Ottonia:* A lineage of Neotropical rainforest shrubs. Systematic Botany 41: 253–262.

Moreira X, Abdala-Roberts L, Galmán A, Francisco M, de la Fuente M, Butrón A, Rasmann S. 2018. Assessing the influence of biogeographical region and phylogenetic history on chemical defences and herbivory in *Quercus* species. Phytochemistry 153: 64–73.

Oksanen J, Blanchet FG, Kindt R, Legendre P, Minchin PR, O’hara RB, Simpson GL, Solymos P, Stevens MH, Wagner H, Oksanen MJ. 2013. Package ‘vegan’. Community Ecology Package, version 2: 1–295.

Orme D, Freckleton R, Thomas G, Petzoldt T, Fritz S, Isaac N, Pearse W. 2018. Caper: comparative analyses of phylogenetics and evolution in R. R package version 1.0.1. https://CRAN.R-project.org/package=caper

Paetzold C, Wood KR, Eaton D, Wagner WL, Appelhans MS. 2019. Phylogeny of Hawaiian *Melicope* (Rutaceae): RAD-Seq resolves species relationships and reveals ancient introgression. Frontiers in Plant Science 10: 1074

Pagel, M. 1994. Detecting correlated evolution on phylogenies: a general method for the comparative analysis of discrete characters. Proceedings of the Royal Society of London. Series B: Biological Sciences 255: 37–45.

Paradis E, Claude J, Strimmer K. 2004. APE: Analyses of Phylogenetics and Evolution in R language. Bioinformatics 20: 289–290.

Parchman TL, Gompert Z, Mudge J, Schilkey F, Benkman CW, Buerkle CA. 2012. Genomewide association genetics of an adaptive trait in lodgepole pine. Molecular Ecology 21: 2991–3005.

Parchman TL, Jahner JP, Uckele KA, Galland LM, Eckert AJ. 2018. RADseq approaches and applications for forest tree genetics. Tree Genetics & Genomes 14: 39.

Parmar VS, Jain SC, Bisht KS, Jain R, Taneja P, Jha A, Tyagi OD, Prasad AK, Wengel J, Olsen CE, Boll PM. 1997. Phytochemistry of the genus *Piper*. Phytochemistry 46: 597–673.

Pavoine S, Ollier S, Pontier D, Chessel D. 2008. Testing for phylogenetic signal in phenotypic traits: new matrices of phylogenetic proximities. Theoretical population biology 73: 79–91.

Peterson BK, Weber JN, Kay EH, Fisher HS, Hoekstra HE. 2012. Double digest RADseq: an inexpensive method for *de novo* SNP discovery and genotyping in model and non-model species. PLoS ONE 7: e37135.

R Core Team. 2020. R: A language and environment for statistical computing. R Foundation for Statistical Computing, Vienna, AT. https://www.R-project.org/.

Rambaut A, Drummond AJ, Xie D, Baele G, Suchard MA. 2018. Posterior summarization in Bayesian phylogenetics using Tracer 1.7. Systematic Biology 67: 901–904.

Rasmann S, Agrawal AA. 2011. Latitudinal patterns in plant defense: evolution of cardenolides, their toxicity and induction following herbivory. Ecology Letters 14: 476–483.

Revell LJ. 2012. phytools: an R package for phylogenetic comparative biology (and other things). Methods in Ecology and Evolution 3: 217–223.

Richards LA, Dyer LA, Forister ML, Smilanich AM, Dodson CD, Leonard MD, Jeffrey CS. 2015. Phytochemical diversity drives plant–insect community diversity. Proceedings of the National Academy of Sciences 112: 10973–10978.

Richards LA, Dyer LA, Smilanich AM, Dodson CD. 2010. Synergistic effects of amides from two *Piper* species on generalist and specialist herbivores. Journal of Chemical Ecology 36: 1105–1113.

Richards LA, Glassmire AE, Ochsenrider KM, Smilanich AM, Dodson CD, Jeffrey CS, Dyer LA. 2016. Phytochemical diversity and synergistic effects on herbivores. Phytochemistry Reviews 15: 1153–1166.

Richards LA, Lampert EC, Bowers MD, Dodson CD, Smilanich AM, Dyer LA. 2012. Synergistic effects of iridoid glycosides on the survival, development and immune response of a specialist caterpillar, *Junonia coenia* (Nymphalidae). Journal of Chemical Ecology 38: 1276–1284.

Richards LA, Oliveira C, Dyer LA. 2018. Shedding light on chemically mediated tri-trophic interactions: A 1h-1H-NMR network approach to identify compound structural features and associated biological activity. Frontiers in Plant Science 9: 1155.

Rognes T, Flouri T, Nichols B, Quince C, Mahé F. 2016. VSEARCH: a versatile open source tool for metagenomics. PeerJ 4: e2584.

Romeo JT, Saunders JA, Barbosa P. 2013. Phytochemical diversity and redundancy in ecological interactions, vol. 30. Berlin, DE: Springer Science & Business Media.

Salazar D, Jaramillo MA, Marquis RJ. 2016. Chemical similarity and local community assembly in the species rich tropical genus *Piper*. Ecology 97: 3176–3183.

Salazar D, Lokvam J, Mesones I, Vásquez P, Zuñiga JMA, de Valpine P, Fine PVA. 2018. Origin and maintenance of chemical diversity in a species-rich tropical tree lineage. Nature Ecology & Evolution 2: 983.

Salehi B, Zakaria ZA, Gyawali R, Ibrahim SA, Rajkovic J, Shinwari ZK, Khan T, Sharifi-Rad J, Ozleyen A, Turkdonmez E, Valussi M. 2019. *Piper* species: a comprehensive review on their phytochemistry, biological activities and applications. Molecules 24:1364.

Sedio BE. 2013. Trait evolution and species coexistence in the hyperdiverse tropical tree genus Psychotria. PhD thesis, University of Michigan, Ann Arbor, MI, USA.

Sedio BE. 2017. Recent breakthroughs in metabolomics promise to reveal the cryptic chemical traits that mediate plant community composition, character evolution and lineage diversification. New Phytologist 214: 952–958.

Sedio BE, Parker JD, McMahon SM, Wright SJ. 2018. Comparative foliar metabolomics of a tropical and a temperate forest community. Ecology 99: 2647–2653.

Sedio BE, Archibold AD, Echeverri JC, Debyser C, Wright SJ. 2019. A comparison of inducible, ontogenetic, and interspecific sources of variation in the foliar metabolome in tropical trees. PeerJ 7: e7536.

Smilanich AM, Dyer LA, Chambers JQ, Bowers MD. 2009. Immunological cost of chemical defence and the evolution of herbivore diet breadth. Ecology Letters 12: 612–621.

Smith JF, Stevens AC, Tepe EJ, Davidson C. 2008. Placing the origin of two species-rich genera in the late cretaceous with later species divergence in the tertiary: a phylogenetic, biogeographic and molecular dating analysis of *Piper* and *Peperomia* (Piperaceae). Plant Systematics and Evolution 275: 9.

Strutzenberger P, Brehm G, Fiedler K. 2012. DNA barcode sequencing from old type specimens as a tool in taxonomy: a case study in the diverse genus Eois (Lepidoptera: Geometridae). PLoS One 7: e49710.

Thompson JN. 1989. Concepts of coevolution. Trends in Ecology & Evolution 4: 179–183.

Thompson JN, Pellmyr O. 1991. Evolution of oviposition behavior and host preference in Lepidoptera. Annual Review of Entomology 36: 65–89.

Wagner CE, Keller I, Wittwer S, Selz OM, Mwaiko S, Freuter L, Sivasundar A, Seehausen O. 2013. Genome-wide RAD sequence data provide unprecedented resolution of species boundaries and relationships in the Lake Victoria cichlid adaptive radiation. Molecular Ecology 22: 787–798.

Wilson J, Forister M, Dyer LA, O’Conner JM, Burls K, Feldman CR, Jaramillo MA, Miller JS, Rodríguez-Castañeda, Tepe EJ, et al. 2012. Host conservatism, host shifts and diversification across three trophic levels in two Neotropical forests. Journal of Evolutionary Biology 25: 532–546.

Wink M. 2003. Evolution of secondary metabolites from an ecological and molecular phylogenetic perspective. Phytochemistry 64: 3–19.

Zagrobelny M, Bak S, Rasmussen AV, Jørgensen B, Naumann CM, Møller BL. 2004. Cyanogenic glucosides and plant–insect interactions. Phytochemistry 65: 293–306.

Yonekura-Sakakibara K, Higashi Y, Nakabayashi R. 2019. The origin and evolution of plant flavonoid metabolism. Frontiers in Plant Science 10: 943.

Zhang Y, Deng T, Sun L, Landis JB, Moore MJ, Wang H, Wang Y, Hao X, Chen J, Li S, Xu M. 2020. Phylogenetic patterns suggest frequent multiple origins of secondary metabolites across the seed plant “tree of life”. National Science Review, in press.

Zheng L, Ives AR, Garland T, Larget BR, Yu Y, Cao K. 2009. New multivariate tests for phylogenetic signal and trait correlations applied to ecophysiological phenotypes of nine *Manglietia* species. Functional Ecology 23: 1059–1069.

