## Supplementary material for "Evidence for both phylogenetic conservatism and lability in the evolution of secondary chemistry in a tropical angiosperm radiation"

**Table S1**. Sampling information for all taxa.

| Species | Country | Collector | Voucher (herbarium) | Clade |
|---|---|---|---|---|
| *P. aduncum* var. *cordulatum* (C.DC.) Yunck. | Brazil | M. Kato | *M. Kato K-1978 (SPF)* | Radula |
| *P. arcteacuminatum* Trel. | Costa Rica | E.J. Tepe | *E.J. Tepe 3534 (CR)* | Radula |
| *P. amphioxys* Trel. | Panama | E.J. Tepe | *E.J. Tepe 4044 (PMA)* | Schilleria |
| *P. armatum* Trel. & Ynck. | Peru | E.J. Tepe | *E.J. Tepe 4394 (USM)* | Radula |
| *P. baezense* Trel. | Ecuador | A.E. Glassmire | *A.E. Glassmire YY1 (CINC)* | Radula |
| *P. barbatum* Kunth | Ecuador | E.J. Tepe | *E.J. Tepe 3005 (QCNE)* | Churumayu |
| *P. cabagranum* C.DC. | Costa Rica | E.J. Tepe | *E.J. Tepe 3531 (CR)* | Schilleria |
| *P. carrilloanum* C.DC. | Panama | E.J. Tepe | *E.J. Tepe 4069 (PMA)* | Schilleria |
| *P. cenocladum* C.DC. | Panama | E.J. Tepe | *E.J. Tepe 3999 (PMA)* | Macrostachys |
| *P. chanchamayanum* Trel. | Peru | E.J. Tepe | *E.J. Tepe 4393 (USM)* | Radula |
| *P. changuinolanum* Trel. | Panama | E.J. Tepe | *E.J. Tepe 3961 (PMA)* | Radula |
| *P. chimonanthifolium* Kunth | Brazil | M. Kato | *M. Kato K-1960 (SPF)* | Radula |
| *P. chrysostachyum* C.DC. | Costa Rica | E.J. Tepe | *E.J. Tepe 3482 (CR)* | Radula |
| *P. colonense* C.DC. (1) | Panama | E.J. Tepe | *E.J. Tepe 4032 (PMA)* | Radula |
| *P. colonense* C.DC. (2) | Costa Rica | E.J. Tepe | *E.J. Tepe 3502 (CR)* | Radula |
| *P. crassinervium* Kunth (1) | Costa Rica | E.J. Tepe | *E.J. Tepe 3463 (CR)* | Churumayu |
| *P. crassinervium* Kunth (2) | Brazil | M. Kato | *M. Kato K-1954 (SPF)* | Churumayu |
| *P. culebranum* C.DC. (1) | Costa Rica | E.J. Tepe | *E.J. Tepe 3413 (CR)* | Radula |
| *P. culebranum* C.DC. (2) | Costa Rica | E.J. Tepe | *E.J. Tepe 3527 (CR)* | Radula |
| *P. culebranum* C.DC. (3) | Panama | E.J. Tepe | *E.J. Tepe 4005 (PMA)* | Radula |
| *P. cyanophyllum* Trel. | Costa Rica | E.J. Tepe | *E.J. Tepe 3427 (CR)* | Peltobryon |
| *P. cyphophyllum* C.DC. | Costa Rica | E.J. Tepe | *E.J. Tepe 3541 (CR)* | Radula |
| *P. disparipes* Trel. (1) | Costa Rica | E.J. Tepe | *E.J. Tepe 3426 (CR)* | Radula |
| *P. disparipes* Trel. (2) | Costa Rica | E.J. Tepe | *E.J. Tepe 3543 (CR)* | Radula |
| *P. disparipes* Trel. (3) | Costa Rica | E.J. Tepe | *E.jJ. Tepe 3486 (CR)* | Radula |
| *P. distigmatum* Yunck. | Panama | E.J. Tepe | *E.J. Tepe 3972 (PMA)* | Isophyllon |
| *P. dryadanum* C.DC. | Panama | E.J. Tepe | *E.J. Tepe 3963 (PMA)* | Radula |
| *P. euryphyllum* C.DC. | Panama | E.J. Tepe | *E.J. Tepe 4000 (PMA)* | Macrostachys |
| *P. figlinum* Trel. | Costa Rica | E.J. Tepe | *E.J. Tepe 3483 (CR)* | Isophyllon |
| *P. fimbriulatum* C.DC. | Panama | E.J. Tepe | *E.J. Tepe 3956 (PMA)* | Macrostachys |
| *P. friedrichsthalii* C.DC. | Costa Rica | E.J. Tepe | *E.J. Tepe 131 (CR)* | Radula |
| *P. gaudichaudianum* Kunth (1) | Brazil | M. Kato | *M. Kato K-1949 (SPF)* | Radula |
| *P. gaudichaudianum* Kunth (2) | Brazil | M. Kato | *M. Kato K-1983 (SPF)* | Radula |
| *P. goesii* Yunck. | Brazil | M. Kato | *M. Kato K-1964 (SPF)* | Schilleria |
| *P. gonocarpum* Trel. | Panama | E.J. Tepe | *E.J. Tepe 3959 (PMA)* | Isophyllon |
| *P. hartwegianum* (Benth.) C.DC. | Panama | E.J. Tepe | *E.J. Tepe 3966 (PMA)* | Macrostachys |
| *P. hispidum* Sw. (1) | Costa Rica | E.J. Tepe | *E.J. Tepe 3430 (CR)* | Radula |
| *P. hispidum* s.l. (2) | Costa Rica | E.J. Tepe | *E.J. Tepe 3476 (CR)* | Radula |
| *P. hispidum* Sw. (3) | Costa Rica | E.J. Tepe | *E.J. Tepe 3496 (CR)* | Radule |
| *P. hispidum* Sw. (4) | Costa Rica | E.J. Tepe | *E.J. Tepe 3509 (CR)* | Radula |

| | | | | |
|---|---|---|---|---|
| *P. hispidum* Sw. (5) | Costa Rica | E.J. Tepe | *E.J. Tepe 3537 (CR)* | Radula |
| *P . holdridgeanum* W.C.Burger | Costa Rica | E.J. Tepe | *E.J. Tepe 4196 (CR)* | Unclassified |
| *P. lagoense* C.DC. | Brazil | M. Kato | *M. Kato K-1944 (SPF)* | Radula |
| *P. latibracteum* C.DC. | Panama | E.J. Tepe | *E.J. Tepe 4050 (PMA)* | Isophyllon |
| *P. longicaudatum* Trel. & Yunck. | Ecuador | E.J. Tepe | *E.J. Tepe 3036 (QCNE)* | Radula |
| *P. lucigaudens* C.DC. (1) | Panama | E.J. Tepe | *E.J. Tepe 3993 (PMA)* | Radula |
| *P. lucigaudens C.DC.* (2) | Panama | E.J. Tepe | *E.J. Tepe 4028 (PMA)* | Radula |
| *P. malacophyllum* (C.Presl.) C.DC. | Brazil | M.Kato | *M. Kato K-1945 (SPF)* | Radula |
| *P. maranyonense* Trel. | Ecuador | E.J. Tepe | *E.J. Tepe 3034 (QCNE)* | Peltobryon |
| *P. mollicomum* Kunth | Brazil | M. Kato | *M. Kato K-1942 (SPF)* | Radula |
| *P. mosenii* C.DC. | Brazil | M. Kato | *M. Kato 1948 (SPF)* | Radula |
| *P. peracuminatum* C.DC. (1) | Costa Rica | E.J. Tepe | *E.J. Tepe 3433 (CR)* | Radula |
| *P. peracuminatum* C.DC. (2) | Panama | E.J. Tepe | *E.J. Tepe 4062 (PMA)* | Radula |
| *P. persubulatum* C.DC. | Panama | E.J. Tepe | *E.J. Tepe 4068 (PMA)* | Radula |
| *P. polytrichum* C.DC. (1) | Panama | E.J. Tepe | *E.J. Tepe 3965 (CR)* | Radula |
| *P. polytrichum C.DC.* (2) | Costa Rica | E.J. Tepe | *E.J. Tepe 3470 (CR)* | Radula |
| *P. pseudofuligineum* C.DC. | Costa Rica | E.J. Tepe | *E.J. Tepe 3499 (CR)* | Radula |
| *P. pseudogaragaranum* Trel. | Panama | E.J. Tepe | *E.J. Tepe 4041 (PMA)* | Radula |
| *P. sancti-felicis* Trel. | Costa Rica | E.J. Tepe | *E.J. Tepe 3415 (CR)* | Radula |
| *P. schuppii* A.H. Gentry | Ecuador | E.J. Tepe | *E.J. Tepe 1562 (QCNE)* | Radula |
| *P. silvivagum* C.DC. | Costa Rica | E.J. Tepe | *E.J. Tepe 3523 (CR)* | Radula |
| *P. sp* | Panama | E.J. Tepe | *E.J. Tepe 3974 (PMA)* | Radula |
| *P. tectoniifolium* Kunth | Brazil | M. Kato | *M. Kato K-1958 (SPF)* | Churumayu |
| *P. tecumense* Trel. | Panama | E.J Tepe | *E.J. Tepe 4008 (PMA)* | Schilleria |
| *P. tuberculatum* Jacq. | Panama | E.J. Tepe | *E.J. Tepe 4039 (PMA)* | Hemipodion |
| *P. umbellatum* L. | Panama | E.J. Tepe | *E.J. Tepe 3967 (PMA)* | Pothomorphe |
| *P. vicosanum* Yunck. | Brazil | M. Kato | *M. Kato K-1966 (SPF)* | Isophyllon |
| *P. villalobosense* Yunck. | Ecuador | E.J. Tepe | *E.J. Tepe 2952 (QCNE)* | Radula |
| *P. villiramulum* C.DC. | Panama | E.J. Tepe | *E.J. Tepe 4031 (PMA)* | Radula |
| *P. xanthostachyum* C.DC. | Costa Rica | E.J. Tepe | *E.J. Tepe 3542 (CR)* | Radula |
| *P. zacatense* C.DC. | Costa Rica | E.J. Tepe | *E.J. Tepe 3438 (CR)* | Radula |

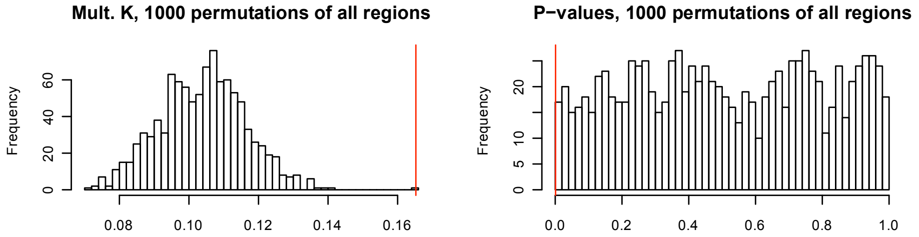


**Figure S1**: Results of the multivariate K test on 1000 permutations of all chemical regions indicate that our observed, significant phylogenetic signal is not an artifact of zero inflation exhibited by the $^{1}$H NMR data. Vertical red lines represent our observed values for the multivariate K statistic and its associate *P*-value.
